# Crystal ribcage: a platform for probing real-time lung function at cellular resolution in health and disease

**DOI:** 10.1101/2022.10.28.514251

**Authors:** Rohin Banerji, Gabrielle N. Grifno, Linzheng Shi, Dylan Smolen, Rob LeBourdais, Johnathan Muhvich, Cate Eberman, Bradley E. Hiller, Jisu Lee, Kathryn Regan, Siyi Zheng, Sue Zhang, John Jiang, Ahmed A. Raslan, Julia C. Breda, Riley Pihl, Katrina Traber, Sarah Mazzilli, Giovanni Ligresti, Joseph P. Mizgerd, Béla Suki, Hadi T. Nia

## Abstract

Understanding the dynamic pathogenesis and treatment response in pulmonary diseases requires probing the lung at cellular resolution in real-time. Despite recent progress in intravital imaging, optical imaging of the lung during active respiration and circulation has remained challenging. Here, we introduce the crystal ribcage: a transparent ribcage that (i) allows truly multiscale optical imaging of the lung in health and disease from whole-organ to single cell, (ii) enables the modulation of lung biophysics and immunity through intravascular, intrapulmonary, intraparenchymal, and optogenetic interventions, and (iii) preserves the 3-D architecture, air-liquid interface, cellular diversity, and respiratory-circulatory functions of the lung. Utilizing these unprecedented capabilities on murine models of primary and metastatic lung tumors, respiratory infection, pulmonary fibrosis, emphysema, and acute lung injury we probed how disease progression remodels the respiratory-circulatory functions at the single alveolus and capillary levels. In cancer, we identified the earliest stage of tumorigenesis that compromises alveolar and capillary functions, a key state with consequences on tumor progression and treatment response. In pneumonia, we mapped mutual links between the recruited immune cells and the alveolar-capillary functions. We found that neutrophil migration is strongly and reversibly responsive to vascular pressure with implications for understanding of how lung physiology, altered by disease and anatomical location, affects immune cell activities. The crystal ribcage and its broad applications presented here will facilitate further studies of real-time remodeling of the alveoli and capillaries during pathogenesis of nearly any pulmonary disease, leading to the identification of new targets for treatment strategies.

## Introduction

The lung is continuously exposed to mechanical, biological, and immunological stresses and is the site of many fatal pathologies such as primary and metastatic cancers, respiratory infections, and both obstructive and restrictive diseases. Our limited understanding of lung pathophysiology was exemplified by the COVID-19 pandemic. Mechanistic understanding of the complex structure-function of the lung at the cellular level and its remodeling at the earliest stages of disease requires characterization of cause-effect relationships at multiple scales down to cellular resolution.

Enclosed by the ribcage and continuously in motion, the functioning lung is a challenging organ to study in real-time at cellular resolution. Clinical imaging modalities (e.g., MRI, CT, and PET) cannot spatiotemporally resolve dynamic alveolar function, capillary flow, and cellular trafficking and motility during respiration. Histological analyses are “snapshot” images of fixed lungs at subcellular resolution without temporal dynamics of the lung microphysiology. Intravital microscopy recently enabled high resolution imaging of microcirculation in real-time^1-5^, which led to major discoveries in lung biology and immunity^2,5-7^. However, these methods (i) compromise the respiratory function of the lung, a defining feature of the pulmonary system with crucial effects on the physics, biology, and immunity of the lung^8-11^, and (ii) can only visualize limited alveoli of the lung through a small imaging window, unable to capture whole-organ heterogeneities or track rare and dynamic events that may span beyond the imaging window.

In a major departure from existing approaches, we introduce the crystal ribcage – a transparent ribcage that enables optical imaging of nearly the entire surface of a functioning mouse lung in real-time and at cellular resolution with preserved respiratory and circulatory functions. We combine *in vivo* benefits – preserved cellular diversity and complex lung architecture – with organ-on-chip model capabilities – optical imaging and high controllability of microphysiology – in the crystal ribcage to perturb the biophysics and immunity of the lung through intrapulmonary, intravascular, and optogenetic interventions while visualizing real-time cellular events. This provides enormous opportunities for understanding cause-effect relationships in the pathogenesis of pulmonary diseases. We utilized the crystal ribcage to probe remodeling of alveolar and capillary functions in mouse models of primary and metastatic lung cancer, bacterial infection, pulmonary fibrosis, emphysema, and acute lung injury. We identified the earliest stage of tumorigenesis when alveolar structure-functions are compromised, probed the dynamic circulatory functions of capillaries remodeled by tumor growth and pneumonia, and demonstrated a dramatic and reversible mechano-responsiveness in immune cell migration in the lung.

## Results

### Crystal ribcage: Development, validation, and capabilities

The crystal ribcage is a transparent, biocompatible platform designed to provide a physiological environment for a functioning lung *ex vivo* that enables truly multiscale optical imaging of the lung in health and disease. The crystal ribcage allows 3-D imaging over the entire surface of the lung (**Fig. 1a**), while uniquely combining the benefits of *in vivo* mouse models (cellular diversity, 3-D lung architecture, breathing and circulation) and *in vitro* organ-on-chip models^12^ (imaging advantages and extensive, precise control over microphysiology) (**Fig. 1b**). We designed the crystal ribcage to mimic the geometry and surface properties of the native ribcage to preserve the *in vivo* physiological conditions of the mouse lung during imaging. Using micro computed tomography (μCT) data of mouse chest cavities, we created 3-D molds of the ribcage that were age and strain specific for mice of interest (in this study: C57/B6, FVB N/J and A/J strains, aged 6-12 weeks). The mold was used in multi-step additive, sacrificial, and thermoforming processes to fabricate the crystal ribcage (**Fig. 1c**, **Extended Data Fig. 1**). We engineered the internal crystal ribcage surface to be hydrophilic, mimicking the native parietal pleura in the mouse chest, so that the lung slides freely along it in its natural configuration (**Supplemental Fig. 1a, Supplemental Video 1-4**). We fabricated the crystal ribcage to be 150 μm thick for imaging with high-powered objectives (e.g., 60x), and confirmed that the ribcage geometry did not cause optical aberrations (**Supplemental Fig. 1b,c**). We fabricated rigid and semi-rigid versions of crystal ribcage using polystyrene and polydimethylsiloxane (PDMS), respectively. Both materials are biocompatible, free of autofluorescence and ubiquitous in biological research. The rigid crystal ribcage allows positive-and negative-pressure ventilation as in mechanically assisted or spontaneous breathing, respectively (**Supplemental Video 1**). The rigid crystal ribcage closely mimics the *in vivo* ribcage in quiet breathing as the majority of the lung deformation is achieved by the diaphragm^13^ (also see **Extended Data Fig. 2a,b**). The semi-deformable PDMS ribcage mimics the macroscale ribcage deformation that occurs during *in vivo* events such as exercise, sighs, and coughs. The PDMS ribcage is also pierceable and self-healing to enable intraparenchymal injections. Finally, we confirmed that the crystal ribcage does not affect alveolar respiratory function by directly comparing ventilation inside the crystal ribcage to ventilation inside the intact mouse ribcage (**Extended Data Fig. 2c**).

**FIGURE 1.**
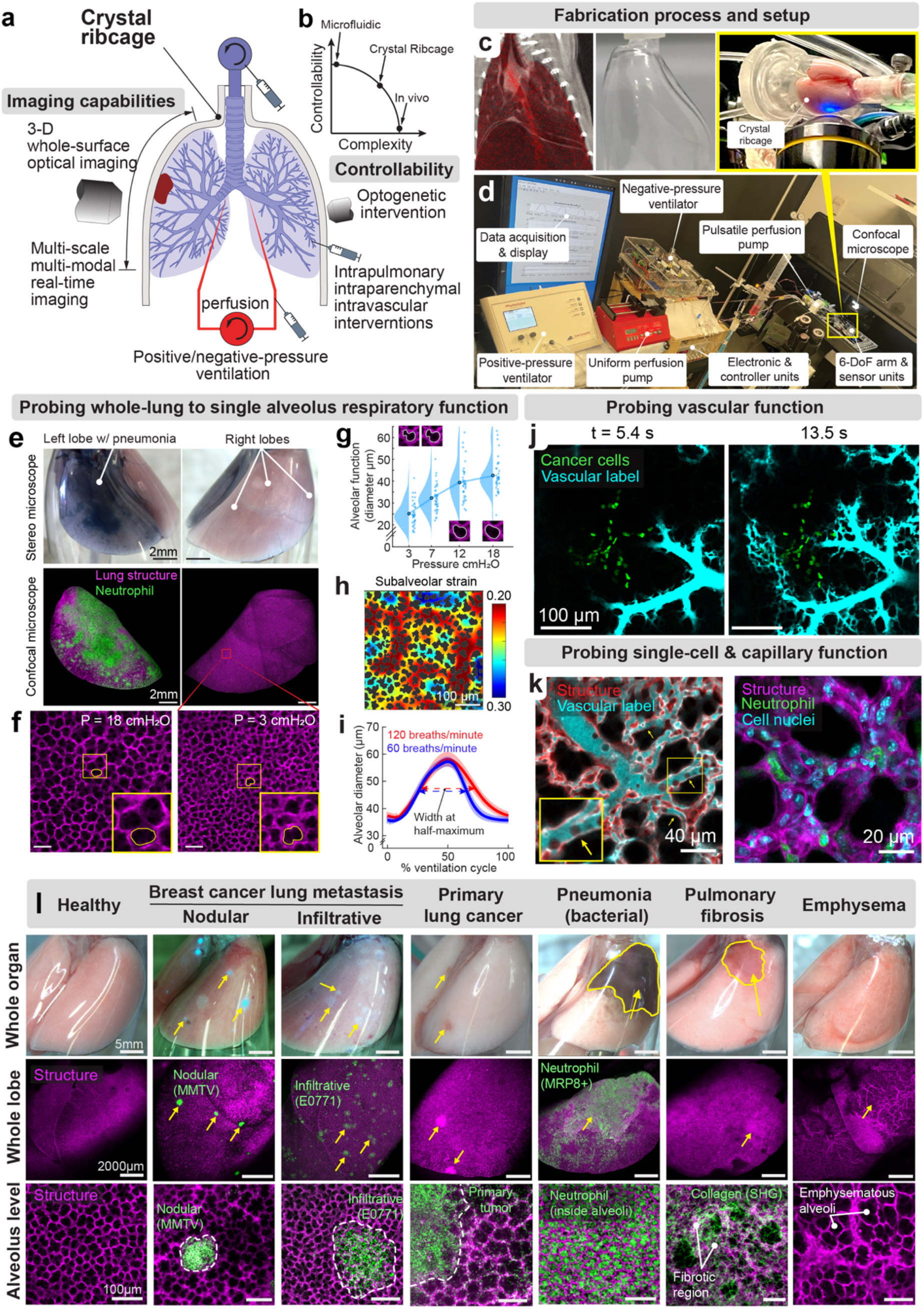
| Development, capabilities, and application of the crystal ribcage in pulmonary research. **(a)** Schematic representation of imaging, controllability, and intervention capabilities of the crystal ribcage. **(b)** The crystal ribcage benefits from the imaging capabilities and controllability of organ-on-chip models while preserving the complexities and cellular diversity of *in vivo* lungs. **(c)** Age- and strain-specific μCT scans used to model the mouse chest cavity to fabricate the crystal ribcage through a multistep additive and sacrificial fabrication process. **(d)** The portable platform to maintain, monitor, and record the lung physiological condition during real-time imaging. **(e)** Imaging the distal lung surface with preserved 3-D geometry, demonstrated by both stereo- and confocal-microscopy imaging of the right and left lobes of a mouse lung with a left-lobar bacterial pneumonia infection (labeled by leakage of intravascular Evans blue into the lung airspace) showing neutrophil infiltration 24 hours after initial infection. **(f)** Imaging desired regions of the lung and tracking alveoli (contoured alveolus for reference) across the respiratory cycle under quasi-static conditions. **(g)** Alveolar function assessed by measuring alveolar diameter as a function of quasi-static alveolar air pressure. The distribution of diameters are connected by the means from m=4 ROI over n=4 mice. **(h)** Representative map of the strain experienced by the alveolar septum during quasi-static ventilation, with sub-alveolus resolution for m=1 ROI. **(i)** Alveolar diameter measured at distinct points of the ventilation cycles at 120 and 60 breaths/min presented as mean ± SEM of data from m=4 ROI over n=2 mice **(j)** Remodeling of single capillary function in lung cancer assessed by real-time confocal microscopy of the spatiotemporal distribution of a Cascade Blue-dextran (10 kDa) flow under quasi-static inflation. **(k)** Cellular activity and alveolar-capillary structure-function imaged during quasi-static inflation, inset yellow box shows a single capillary with shadow of cells inside it. (**l**) The crystal ribcage capabilities demonstrated on models of breast cancer lung metastasis, primary lung cancer, respiratory infection, pulmonary fibrosis, and emphysema from whole organ down to alveolus, capillary, and cell length and functional scales, imaged with confocal and multi-photon microscopy.

The crystal ribcage is equipped with custom and commercial ventilators for negative- and positive-pressure ventilation, both uniform and pulsatile perfusion pumps, and sensors with a data acquisition unit to preserve and assess the lung’s circulatory and respiratory functions (**Fig. 1d**). The crystal ribcage is attached to a custom 6-degree-of-freedom arm to position any region of the lung surface on top of or under a microscope to image the whole lobe or single alveoli using any optical imaging modality (**Fig. 1e**, **Extended Data Fig. 3**). This allows probing anatomically localized pulmonary diseases, such as lobar pneumonia (**Fig. 1e**) and capturing sporadic features such as tumor metastases (**Fig. 1f)**.

We quantified alveolar respiratory function by imaging a specific region of interest (ROI) on the lung surface over the normal inspiratory quasi-static pressure range from 3 to 18 cmH_2_O^14^ (**Fig. 1g**). We then computed the sub-alveolar strain (normalized deformation) field (**Fig. 1h**) over this pressure range, as an indirect measure of alveolar functionality. Next, we studied alveolar mechanics during dynamic positive-pressure ventilation at 120 and 60 breaths/min while maintaining a positive end expiratory pressure (PEEP) of 4 cmH_2_O and a peak inspiratory pressure (PIP) of 12-14 cmH_2_O (**Fig. 1i**, **Extended Data Fig. 4**). We found that alveoli near the base of the lung have a larger diameter for a longer duration of the ventilation cycle at higher respiratory rates (120 breaths/min).

We next probed pulmonary circulation in the context of cancer and other pathologies using the crystal ribcage. First, we imaged the spatiotemporal dynamics of a fluorescent dye distribution (**Fig. 1j**). Second, we imaged single capillary and trafficking of single immune cell within the capillary at high spatiotemporal resolution (**Fig. 1k**, **Supplemental Fig. 2, Supplemental Video 5**). The crystal ribcage simultaneously enables imaging of key resident cell types in the lung, as demonstrated for alveolar type II cells (SPC-GFP+; **Extended Data Fig. 5**) and can be used to visualize the lung illuminated with fluorescent nuclear, membrane, and vascular labels, or through label-free approaches such as second harmonic generation (SHG).

We demonstrated the versatility of the crystal ribcage to probe lung dysfunction in nearly any pulmonary disease or condition with parenchymal presentation (**Fig. 1l**), such as primary and metastatic breast cancer (**Fig. 2**), pneumonia and acute injury (**Fig. 3-5**), emphysema, and fibrosis (**Supplemental Figs. 2,3)** at the whole organ, whole lobe, and alveolar scales.

**FIGURE 2.**
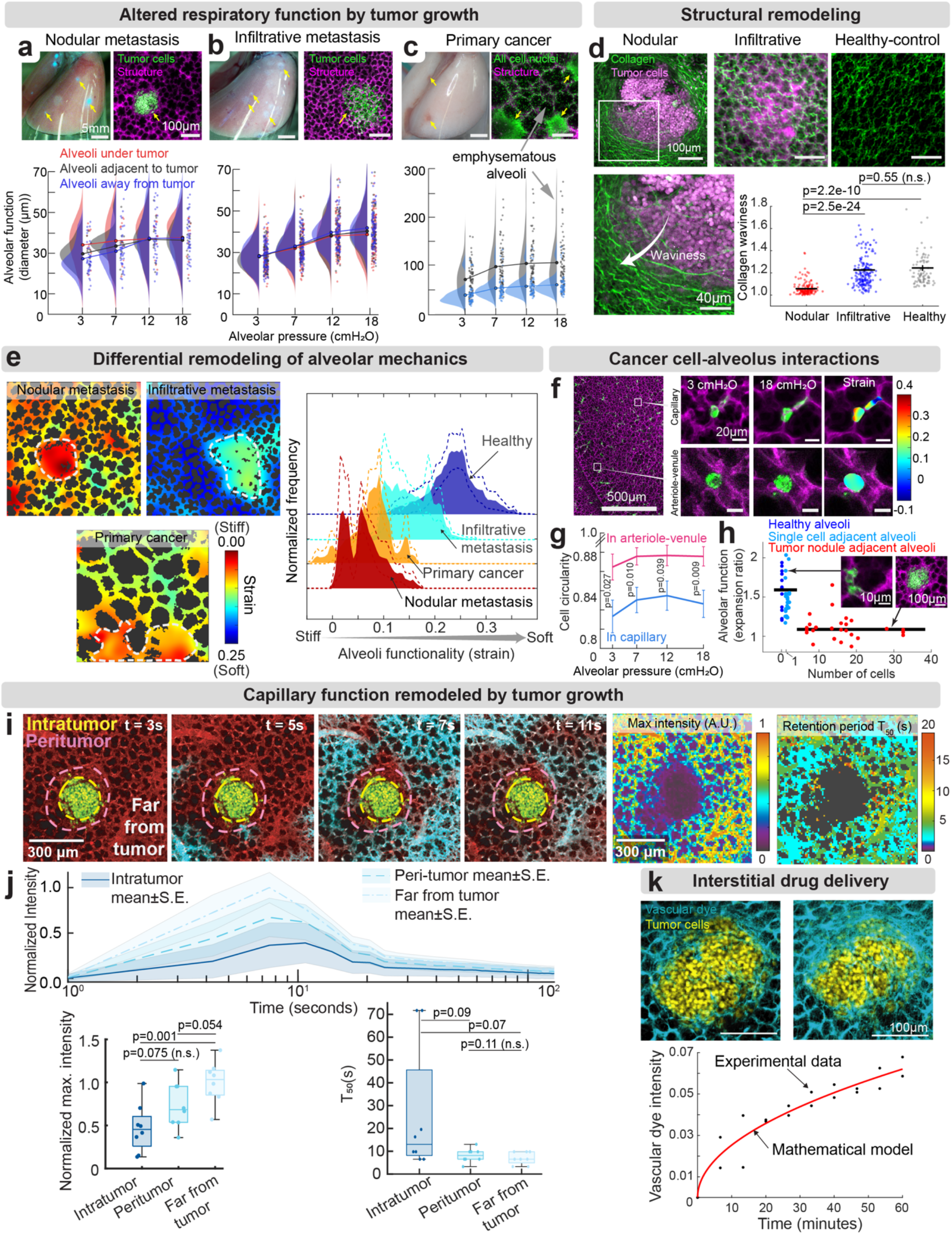
| Multi-scale structure-function changes in lung cancer models. Tumors (yellow arrow) at whole-organ and alveolar scale, distribution of alveolar diameters measured intratumorally, peritumorally, and far from tumor at each quasi-static pressure, line connects mean of distribution **(a)** representative nodular metastasis (green) and alveoli (magenta), for m=4 ROI over n=3 mice, **(b)** representative infiltrative metastasis (green) and nearly fully functional alveoli (magenta), for m=4 ROI over n=3 mice, and **(c)** representative primary lung cancer nodules with adjacent emphysematous alveoli, for m=3 ROI over n=3 mice. **(d)** Second harmonic generation imaging of collagen architecture in nodular (m=7 ROI over n=3 mice) and infiltrative (m=7 ROI over n=2 mice) metastatic cancers, and healthy (m=4 ROI over n=1 mouse), mean line of the scatter data from k=150, 188, 92 fibers respectively. **(e)** Representative alveolar strain as lung pressure increases from 7 to 12 cmH2O in each disease and mean ± SEM strain distribution per model for same ROI and mouse count in **a, b, c. (f)** Representative strain map of arrested cancer cells during quasi-static inflation in capillaries vs arterioles/venules. **(g)**. Bulk comparison of circularity of cancer cells tracked at different pressures taken from n=30 cells in arterioles/venules and n=40 cells in capillaries, mean ± SEM of data is presented, and significance is tested between cells in each group at each pressure. (**h**) Alveolar function decreases with increasing cells in cancer nodule. Individual data points represent nodules and lines are mean alveolar expansion ratio for nodules =< 1 cell and >2 cells from m=60 nodules over n=16 mice. **(i)** Real-time imaging of the disrupted capillary function through pressure-controlled perfusion of fluorescent Cascade Blue-dextran (10kDa), and **(j)** distribution and retention of the same for m=4 ROI over n=5 healthy mice and m=8 ROI over n=4 tumor bearing mice. **(k)** Passive diffusion of a small molecule tracer into the tumor bulk and the mathematically estimated diffusion coefficient (see Methods) of the tracer in the tumor. Boxplots present median with 25th and 75th percentiles, whiskers are the maximum and minimum data points not considered outliers. Comparison between groups performed with two-tailed Student’s t-test for significance<0.05.

### Disruption of alveolar and capillary dynamics by tumor growth

Primary lung cancer is among the leading fatal cancers ^15^. The lung is also a major site of metastasis for many cancer types including breast cancer^16^. While early steps of cancer cell extravasation and immune interactions have been studied using intravital microscopy in a lung immobilized by glue or vacuum^1,3^, understanding of tumor dynamics during active respiration and circulation is lacking.

To answer long-standing questions surrounding remodeling of alveolar structure and function by tumors, we investigated two mouse models of breast cancer lung metastasis with differential growth patterns. According to our previous work^17^ and others^18^, tumor growth patterns are generally categorized as *nodular* (cells grow as a dense nodule pushing away normal tissue) or *infiltrative* (cancer cells replace normal cells with minimal remodeling of existing tissue; also known as co-optive^18^), with distinct biological and clinical outcomes. Infiltrative tumors co-opt the tissue’s existing blood vessels and structure and initiate less angiogenesis^19^. In contrast, nodular tumors disrupt existing microarchitecture and form new blood vessels^18^. We have previously characterized differential growth patterns in infiltrative primary brain tumors (e.g., glioblastoma) versus nodular metastatic brain tumors^17^. Surprisingly, the same two phenotypes were also observed in breast cancer metastasis to the lung. Next, we probed the differential remodeling of the peritumor microenvironment^20^.

We modeled nodular and infiltrative metastatic tumors using murine breast cancer cell lines MCa-M3C-H2B-dendra2 and E0771-H2B-dendra2, respectively (**Fig. 2a,b**). The nodular phenotype tumors appeared to both “push” into the lung tissue and bulge out of the pleural surface of the lung, while the infiltrative phenotype appeared to replace the existing lung tissue and was located superficially on the pleural surface (**Extended Data Fig. 6a,b**). Upon probing the mechanics of local alveoli, we observed that alveoli filled with nodular tumors in the range of 100-200 µm in diameter lacked the ability to change volume (**Fig. 2a**) impairing respiratory function. These alveoli remained maximally dilated across all alveolar pressures, indicating that nodular cancer cells distend and fill the alveolar airspace. Interestingly, functional impairment extended up to 1-2 alveoli into the peritumoral region. Larger 1-2 mm diameter nodules caused alveoli to become stretched and compacted up to 200 µm away from the tumor boundary (**Supplemental Fig. 4,5**, **Supplemental Video 6**). In contrast, the infiltrative tumors did not compromise alveolar respiratory function intratumorally or peritumorally (**Fig. 2b**), suggesting that they merely take over existing tissue structures. In a carcinogen-induced model of primary lung cancer, early-stage tumors presented with a nodular-like phenotype. However, the peritumoral alveoli became emphysematous, as multiple alveoli coalesced into a single large airspace near the lesions (**Fig. 2c**). These emphysema-like airspaces were almost twice as large as alveoli far from the tumor and not as responsive to quasi-static pressure changes (**Fig. 2c**). This indicates that, despite not being directly filled with cancer cells, these alveoli still had reduced functionality compared to unaffected alveoli in the same lung.

Structural remodeling of the extracellular matrix (ECM) by tumors is strongly correlated to progression and invasion of tumors^21,22^. We investigated the collagen structure surrounding nodular and infiltrative metastatic tumors using two-photon second-harmonic generation (SHG) microscopy. Established nodules remodeled the pleural collagen to become stretched in comparison to the wavy collagen structures in the healthy lung (**Fig. 2d**). These straightened fibers indicate that alveolar walls adjacent to nodular tumors are under tension, unable to expand or contract during breathing (**Supplemental Fig. 4h**), reducing local gas exchange. In contrast, collagen fibers surrounding infiltrative tumors retained the same characteristic waviness seen in healthy tissue, implying that the structure-function of the alveoli remained unaffected. We also assessed alveolar function by computing the strain in the walls of single alveoli using a deformable image registration algorithm we adapted and validated for lung microscopy (**Supplemental Fig. 6**). Reduced strain in the septal walls during quasi-static ventilation indicates reduced alveolar functionality. In comparing the strain maps between the three different tumor types, nodular metastatic breast cancer caused the most severe dysfunction followed by the primary cancer nodules in the surrounding alveoli, and infiltrative metastatic breast cancer barely impacted local alveolar strain (**Fig. 2e**).

Next, we studied bi-directional mechanical interactions between metastatic cancer cells and the lung tissue. We first found that single cancer cells sequestered at the capillaries do not impact alveolar strain, but instead experience substantial strain themselves (**Fig. 2f**, **Extended Data Fig. 7**) and have lower circularity at all alveolar pressures in comparison to cells located at larger vessels (**Fig. 2g**). It was previously suggested that cancer cells experiencing high, cyclic mechanical forces contribute to tumorigenesis in *in vitro*^23^. We directly show that single cancer cells within the complex 3-D vascular architecture experience deformation in response to alveolar pressure in a functional lung, similar to what we previously reported for cell-sized polyacrylamide beads of uniform stiffness^24^. We next found metastatic nodules as small as 7-10 cells in diameter (approximately the size of one alveolus) was the critical size needed to reduce local alveolar function due to remodeling by the cancer bulk^25-28^ (**Fig. 2h**).

Next, based on dramatic remodeling of alveoli by nodular tumors, we investigated the remodeled intra- and peri-tumoral capillary function. We perfused a bolus of fluorescent Cascade blue dextran (10 kDa) with RPMI cell culture media (or whole blood, **Supplemental Fig. 7**) under pressure-controlled flow of 15 cmH_2_O (approximate median vascular pressure at the mouse pulmonary artery^29^), creating a flow of ∼1 ml/min into the lung inside the crystal ribcage. We imaged the exclusion of the dye inside the tumor bulk and up to ∼100 µm away from it in the peritumoral region indicating loss of circulatory functions in blood vessels remodeled by metastatic nodules (**Fig. 2i**). The fluorescence intensity was lower, and it took longer for the dye to reach inside the tumor bulk compared to the peritumor and unaffected capillaries far from the nodule (**Fig. 2j**). We also observed similar disruption of capillary function in the presence of dispersed single cancer cells before the tumors were established (**Extended Data Fig. 8, Supplemental Video 7**). We investigated the rate of passive diffusion in established nodular lung tumors, as convective flow was insufficient. Smaller molecular weight fluorescent CBhydrazide tracer (550 Da), was controllably perfused until background fluorescence reached steady state (**Fig. 2k**). By fitting a mathematical model based on the diffusion equation to the rate of increase in fluorescence intensity inside the tumor (see Methods), we estimated the diffusion coefficient of the tracer in nodular tumors, a key parameter determining the rate of the drug and nutrient delivery to the tumor core, to be 3.25 µm^2^/s, consistent with what was reported in melanoma tumors^30^.

### Respiratory and circulatory functions disrupted in pneumonia

Pneumonia has been a leading global burden of healthcare^31^, even prior to the global COVID-19 crisis. Therefore, as another application of crystal ribcage, we decided to probe the impact of bacterial pneumonia infection and acute injury on lung respiratory and circulatory functions from the whole lobe down to single alveolus and single capillary scales. Imaging the whole lung showed that the hepatized left apex, was densely packed with MRP8+ neutrophils both at the whole lobe and alveolar scales (**Fig. 3a,b**). We examined the functionality of alveoli for quasi-static applied alveolar pressures (3-18 cmH_2_O) similar to previous whole-lung pressure-volume curves characterized in mice^14,32,33^. The pressure-volume curves showed substantial heterogeneities (**Fig. 3**). Further analysis revealed that pneumonia-infected alveoli largely remained constant and were similar in size to healthy alveoli imaged at an alveolar pressure of 7 cmH_2_O (**Fig. 3c**). Thus, infected alveoli are unable to respond (expand and contract) to pressure applied at the trachea due to both neutrophil recruitment and liquid influx into the airspace (pulmonary edema). We independently visualized neutrophil recruitment and pulmonary edema in infected alveoli simultaneously at the whole-lobe and single alveolus level (**Supplemental Fig. 8**) and found regions with high neutrophil recruitment closely match pulmonary edema (**Extended Data Fig. 9**). In comparison to disease-free alveoli, the inflammatory response to infection causes a decrease in strain at the lobar (**Supplementary Fig. 9a**) and alveolar scales (**Fig. 3d,e**) due partially to filling of the alveoli with incompressible liquid that increases local surface tension.

**FIGURE 3.**
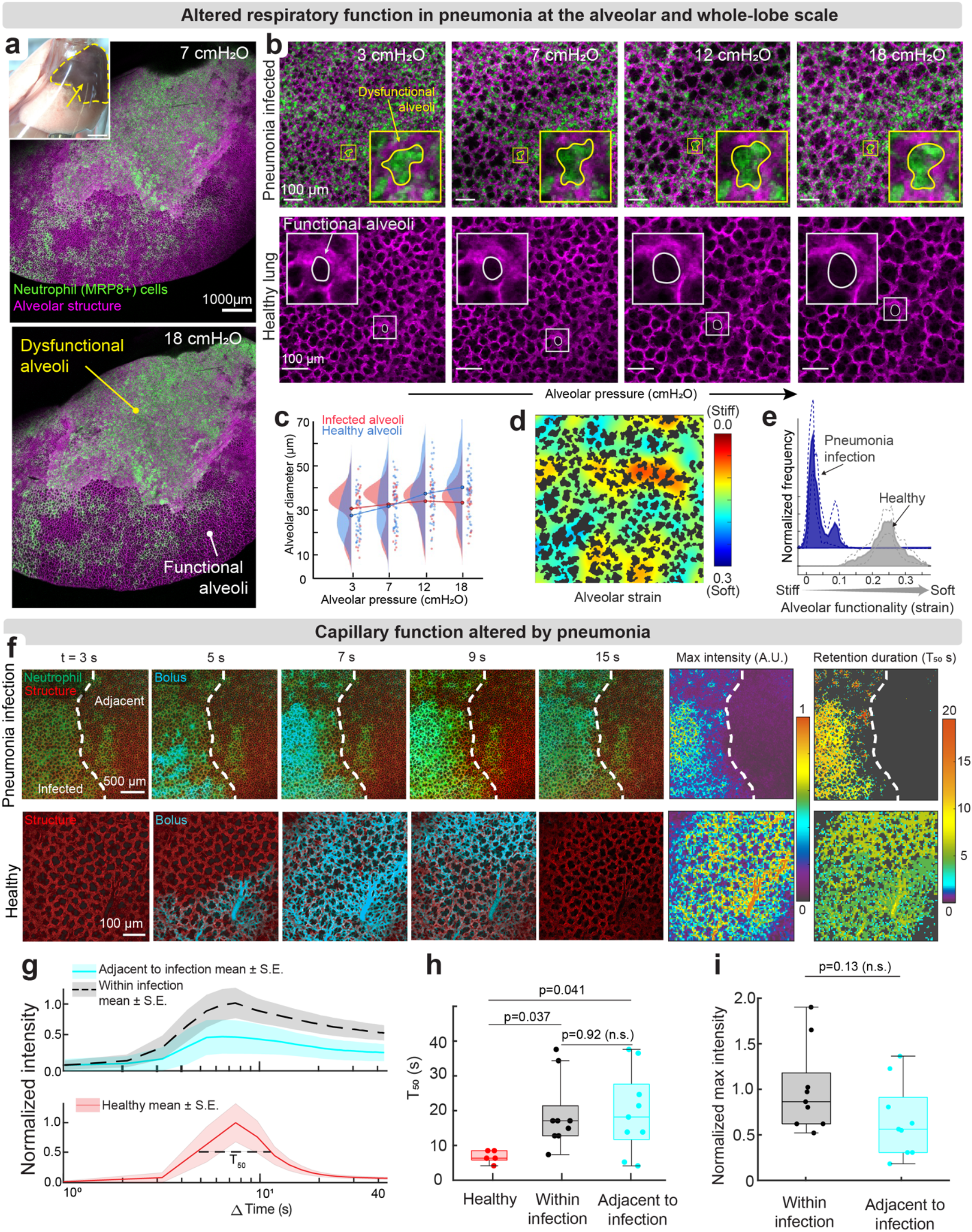
| Impairment of alveolar respiratory and circulatory functions in pneumonia. (**a**) Entire left lobe imaged via confocal microscopy at 7 and 18 cmH2O quasi-static alveolar pressures. (**b**). Representative images of alveolar deformation in a pneumonia-infected versus healthy lung in response to alveolar pressure changes. Single alveolus contoured shows dysfunctional alveoli (marked by neutrophil infiltration) are unresponsive to applied increases in air pressure compared to functional alveoli. (**c**) Distributed alveolar diameter data at each quasi-static pressure connected by the mean line for each infected and healthy groups for m=6 ROI over n=4 mice. (**d**) Representative strain map at alveolar scale strain showing reduced strain in neutrophil recruited areas. (**e**) Mean ± SEM of strain distribution in pneumonia infected and healthy regions, same mouse counts as **c**. (**f**) Capillary function in pneumonia infected and healthy lung perfused with CBhydrazide (550 Da) in RPMI medium at 15 cmH2O pressure-controlled flow and an alveolar pressure of 7 cmH2O. (**g**) Time course of fluorescence intensity in infected, infection-adjacent, and healthy capillaries. (**h**) Time duration when fluorescence intensity in the capillary bed was greater than 50% of the maximum intensity, indicating the tendency for dye to be retained in the capillary bed for m=4 ROI over n=5 healthy mice and m=9 ROI over n=4 pneumonia-infected mice. (**i**) Maximum fluorescence intensity in healthy and infection-adjacent capillaries after delivery of the bolus, for same mouse count as **h**. Boxplots present median with 25th and 75th percentiles, whiskers are the maximum and minimum data points not considered outliers. Comparison between groups performed with two-tailed Student’s t-test for significance < 0.05.

We next asked whether the recruitment of immune cells and edema in pneumonia affects capillary function and perfused a bolus of small molecule CBhydrazide (550 Da) fluorescent tracer to simulate a systemic drug delivered to treat infections. In real-time we mapped the heterogenous distribution (**Fig. 3f**, **Supplemental Video 8**) and intravascular delivery (**Fig. 3g,h**) of the perfusate near the infected regions, representing the retention and clearance of the drug. There was no difference in tracer arrival but delayed clearance from the infected region (**Fig. 3f,g**), as the duration of high fluorescence in pneumonia infected tissue was greater than adjacent healthy tissue

(**Fig. 3h**). This indicates that local vascular function is likely compromised in the injured alveoli, causing the fluorescent tracer to leak into the interstitial space from the vascular space such that it cannot be easily cleared by continuous perfusion. Additionally, vascular dynamics were slower in the injured region in comparison to healthy tissue, suggesting that the dye was likely transported by diffusion rather than convection in the injured tissue (**Fig 3f-i**). While it is unlikely that the infected tissue is totally isolated from the rest of the pulmonary circulation, given that both healthy and injured tissues experienced a comparable maximum fluorescence intensity upon the initial arrival of the fluorescent tracer in the vasculature (**Fig. 3i**), the leakiness of vessels in pneumonia potentially allowed the dye to be delivered faster than in metastatic tumors (**Supplementary Fig. 9b**).

### Neutrophils are reversibly mechano-responsive

While we observed that recruitment of immune cells alters alveolar and capillary functions, we endeavored to see if the relationship was mutual, i.e., whether immune cell activity is also sensitive to changes in alveolar or capillary pressure. Neutrophil recruitment from pulmonary capillaries has differing rules, mechanisms, and constraints than from the systemic post-capillary venules^34-37^. Particularly, neutrophil emigration in the pneumonic lung is primarily from pulmonary capillaries rather than post capillary venules, as is characteristic of other tissues. Intravital microscopy has previously visualized neutrophil migration in the lung through implanted windows on the mouse ribcage^37,38^. However, these approaches cannot controllably alter alveolar and capillary pressures. We tracked single neutrophils in real-time while modulating alveolar and vascular pressures to probe the effects of breathing and circulation on pulmonary immunity.

We observed dramatic and reversible changes in neutrophil migration speed in response to an applied increase in vascular pressure in mouse models of lipopolysaccharide (LPS) injury and in bacterial pneumonia. It has been reported that the mean arterial pressure in the catheterized *in situ* resting mouse is 13-20 cmH_2_O^39-41^, and the left atrial pressure is approximately 5.4 cmH_2_O^42^. We chose to apply a constant vascular pressure across the capillary bed of up to 15 cmH_2_O^43,44^ as an approximation of the pressure drop between the pulmonary artery and left atrium. In a lobar LPS model of acute injury in MRP8-mTmG mice^45,46^ (**Fig. 4a**), we tracked neutrophils migrating within the injured alveoli in real-time and found that neutrophils migrated longer distances (**Fig. 4b**) and nearly twice as fast (**Fig. 4c**) at 15 cmH_2_O compared to zero vascular pressure. This change in speed was also reversible, as neutrophil speed responded to repeated step-changes in vascular pressure (**Supplemental Video 9, Supplemental Fig. 10**). We maintained zero-flow conditions to decouple the effects of vascular flow and shear forces on immune cells^47^. We also measured the persistence (ratio of end-to-end displacement to distance travelled) and found that it did not increase with increasing vascular pressure in the lung implying the increase in cell motility is not flow-driven (**Fig. 4d**). Given no vascular flow and that neutrophil emigration was agnostic to pressure changes, it is unlikely that soluble chemokines (the majority of which are bound to proteoglycans) redistribute quickly enough to play a major role in our observed changes in migration speed. Additionally, the geometric and biophysical constraints posed by the lung capillaries largely prevent neutrophil rolling along endothelial cells, meaning that selectin binding is an unlikely to be a strong contributor to the differences in migration speed we observe here^34-37^. Our results suggest that, within the complex environment of the lung capillary bed, neutrophils either directly respond to changes in vascular pressure or they are indirectly influenced by other factors that change with vascular pressure, such as vascular diameter, the surface density of adhesion molecules and chemokines on the luminal surface of endothelial cells, or alteration in cell junctions in response to stretch^48-51^.

**FIGURE 4.**
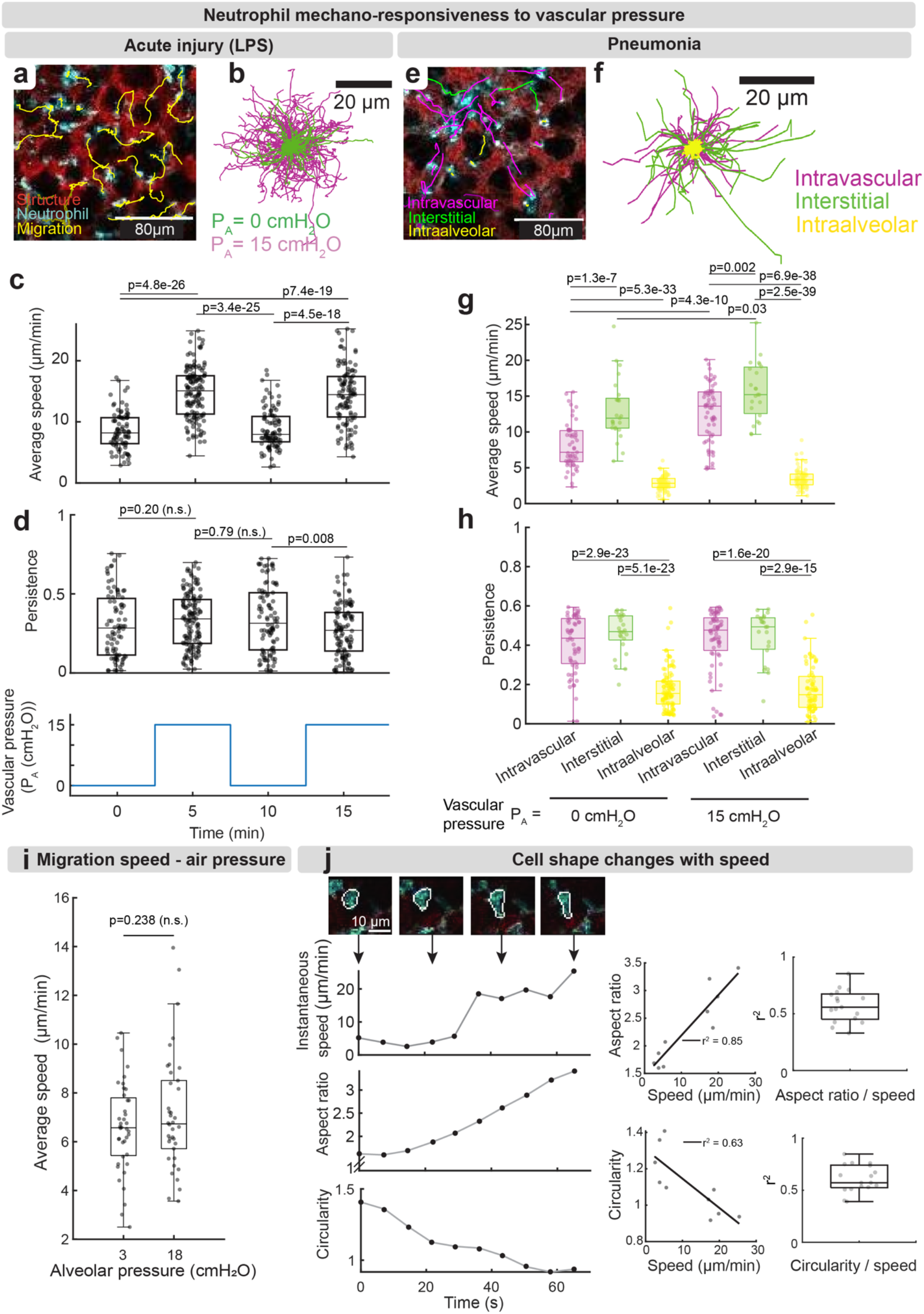
| Neutrophil migration is reversibly mechano-responsive to vascular pressure. (**a**) Representative image of neutrophil migration with cell trajectories (yellow) in the LPS model of acute lung injury. (**b**) Neutrophil migration map of paths travelled by single neutrophils over a 5-minute imaging interval. (**c**) Neutrophil average speed increases with vascular pressure and is reversible in LPS-damaged lungs (n=3 mice), for a constant alveolar pressure of 7 cmH2O. (**d**) Persistence directionality of neutrophil migration in LPS-damaged lungs (n=3 mice). (**e**) Representative traced neutrophil migration in intra-alveolar (yellow), interstitial (green), and intravascular (magenta) spaces in Sp3 pneumonia lungs at a pulmonary artery pressure (PA) of 15 cmH2O and a constant alveolar pressure of 7 cmH2O. (**f**) Neutrophil migration map for 15 cmH2O hydrostatic perfusion pressures and a constant alveolar pressure of 7 cmH2O in pneumonia lungs for 5 minutes. (**g**) Neutrophil average speed and (**h**) persistence in intra-alveolar, interstitial, and intravascular spaces under 0 or 15 cmH2O hydrostatic perfusion pressures in pneumonia lungs (n=3 mice), for a constant alveolar pressure of 7 cmH2O. (**i**) Neutrophil average speed in pneumonia lungs under quasi-static alveolar pressures of 3 (m=38 neutrophil across n=2 mice) versus 18 cmH2O (m=40 neutrophil across n=2 mice). (**j**) Representative changes in single neutrophil speed, aspect ratio, circularity of the same neutrophil with its respective aspect ratio vs. speed and circularity vs. speed linear regressions, and the regression coefficient of determination of aspect ratio vs. speed and circularity vs. speed of a population of neutrophils (m=27 cells over n=3 mice). Boxplots present median with 25th and 75th percentiles, whiskers are the maximum and minimum data points not considered outliers. Comparison between groups performed with two-tailed Student’s t-test for significance < 0.05.

We independently confirmed the mechano-responsive behavior of neutrophils in bacterial pneumonia (Sp3) infection. Imaging migration of neutrophils around infected alveoli (**Fig. 4e,f**) revealed that cells located in the vascular and interstitial space migrated farther compared to neutrophil inside the alveoli. Additionally, neutrophils in the vasculature and interstitium moved almost twice as fast when vascular pressure was increased from 0 to 15 cmH_2_O, while intra-alveolar neutrophils remained slow regardless of vascular pressure changes (**Fig. 4g**). The persistence of neutrophils in the pneumonia model was also independent of changes in vascular pressure (**Fig. 4h**). We concurrently found that neutrophil speed did not respond to alveolar pressure (**Fig. 4i**). Lastly, we observed that neutrophils dramatically changed shape while migrating in lung capillaries and became more elliptical when moving faster (**Fig. 4j**, **Supplemental Video 10**). We utilized the combined benefits of high controllability and intact cellular diversity of the complex vascular architecture in the crystal ribcage to probe the mechano-immunity of neutrophils, which potentially extends to other immune cell types^52^.

### Respiration-circulation coupling is disrupted in lung injury

Lung respiration and circulation dynamics are strongly coupled. At the whole organ scale, this regulates the diameter of large blood vessels and in turn determines pulmonary vascular resistance^53^. To our knowledge respiration-circulation coupling has not been characterized at the alveolar-capillary scale. Hence, we quantified the effect of respiration-circulation coupling on capillary and arteriole/venule distension in healthy lungs by quasi-statically varying physiologically relevant alveolar and vascular pressure (**Fig. 5a**, **Supplemental Fig. 11**). We found the vascular diameter (inclusive of wall thickness and lumen width) in arterioles-venules to first increase and then decrease, creating a “bell shaped” curve across the applied vascular pressure range when alveolar pressure increased over the inspiratory range from 3 to 18 cmH_2_O. This pressure response implies arteriole/venule diameter is maximized, resulting in a minimum vascular resistance, for an optimal range of alveolar pressures (7-12 cmH_2_O) (**Fig. 5b**). Capillary diameter increased dramatically with increasing vascular pressure (**Fig. 5c**), but it started monotonically decreasing for alveolar pressures higher than 7 cmH_2_O, for intermediate vascular pressures only (**Fig. 5c**). This indicated that alveolar pressures near 7 cmH_2_O minimize vascular resistance in capillaries.

**FIGURE 5.**
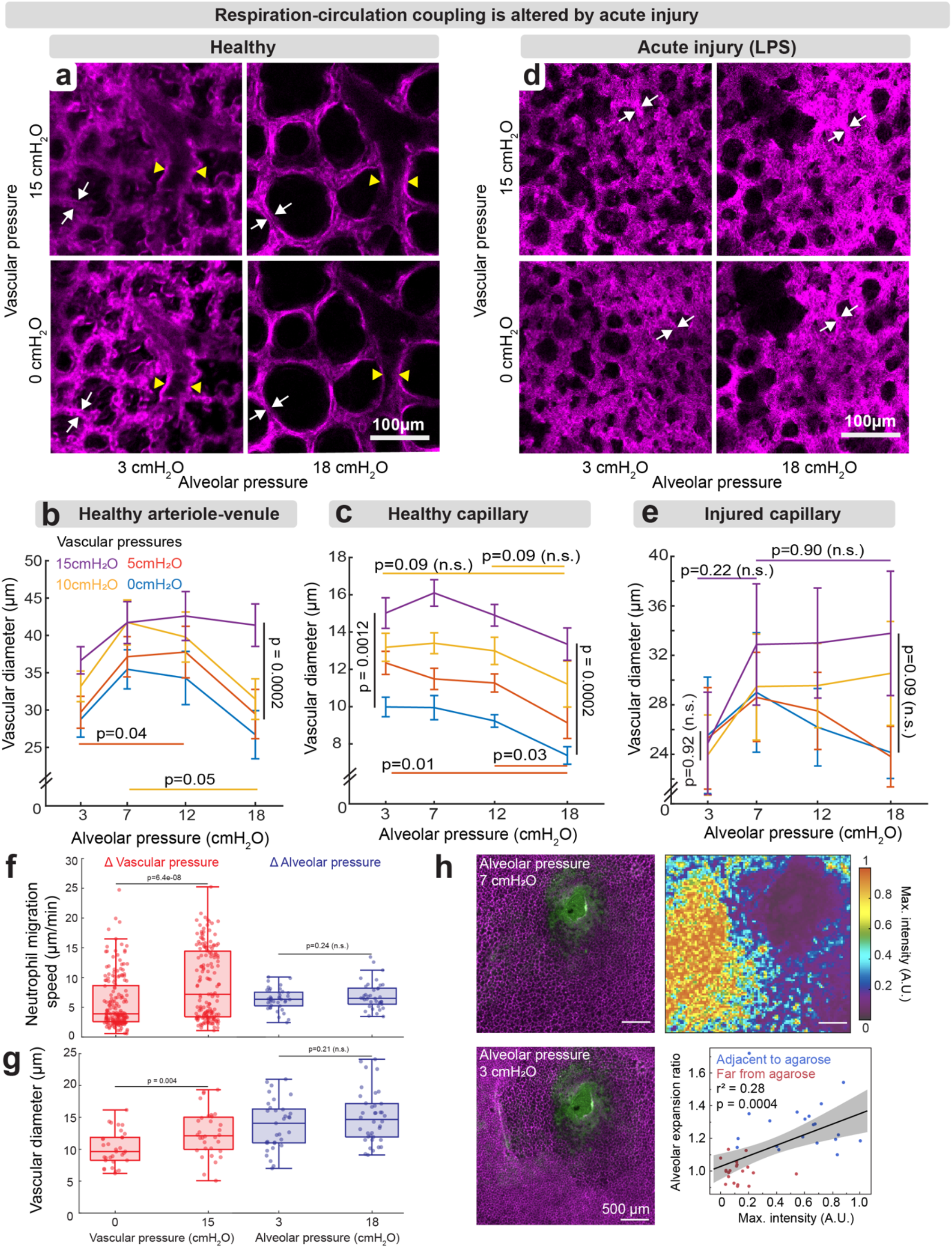
| Respiration-circulation coupling in healthy lungs and its disruption by acute injury. (**a**) Representative image of structural changes in the lung associated with variable quasi-static alveolar and vascular pressures, with measurements of diameter changes in single lung capillaries (white arrows) and large arterioles (yellow arrows), which can be tracked in the crystal ribcage across normal respiration-circulation parameters. Respiration-circulation coupling trends can be observed by measuring the vascular diameter in response to changing pressures in (**b**) large, heathy arterioles and venules (m=15 ROI over n=3 mice) and (**c)** healthy capillaries (m=19 ROI over n=4 mice). (**d**) Representative images of the injured lung after challenge with LPS, showing that single capillaries (white arrows) affected by acute injury (LPS) become inflamed and distend differently for the same pressure range. (**e**) Capillaries acutely injured after LPS challenge become non-responsive to changes in alveolar and vascular pressure (m=16 ROI over n=4 mice). All traces are presented as mean ± SEM and the p-value determined by testing groups using two-tailed Student’s t-test for significance < 0.05. (**f**) Neutrophils migrate at faster speeds as vascular pressure measured at the pulmonary artery (PA) increases (m>130 neutrophils over n=2 mice) compared with little effect of increasing air pressure in the alveoli (m=40 neutrophils over n=2 mice). (**g**) Similarly vascular diameter changes are significantly different with increasing vascular pressure (n=2 mice), while changing alveolar pressure has little effect on it (n=2 mice). (**h**) Injecting Evans blue labeled agarose (green) produces a local blockage in the lung that disrupts alveolar-capillary function, as alveoli near the blockage do not deform across quasi-static air pressures and exclude perfused fluorescent tracer, while alveoli and capillaries farther away from the blockage remain functional (m=40 alveoli over n=1 mouse).

We contrasted the above findings with an acutely injured lung (LPS-treated) in which the parenchyma was substantially remodeled by an inflamed capillary bed and reduced alveolar size across the full breathing cycle (**Fig. 5d**). Respiration-circulation coupling is lost in injured capillaries, as increasing vascular pressure did little to increase vascular diameter. Increasing air pressure also did not reduce vascular diameter, compared to healthy capillaries (**Fig. 5e**). This disrupted the ability of the lung to respond to alveolar and blood pressure changes likely reduce local flow with loss of oxygen transport. The altered respiration-circulation coupling described above may also provide insight into neutrophils’ mechano-responsiveness to pressure. We observed faster migration with increasing vascular pressure in neutrophil populations recruited to a bacterial pneumonia infected lung (**Fig. 5f**), which was accompanied by an increase in capillary diameter (**Fig. 5g**). However, increasing alveolar pressure in pneumonia-injured lungs did not alter neutrophil speeds (**Fig. 5f**), nor did it change capillary diameters (**Fig. 5g**). This correlation may suggest that tensional stresses on the luminal surface of the vasculature, due to increasing vascular diameter, alters neutrophil migration speed by, potentially, facilitating binding of neutrophils to the endothelial wall^54^.

Lastly, we acutely perturbed the lung by injecting solid phase agarose (see Methods) in the parenchyma to disrupt the respiration-circulation coupling. We found that alveoli near the agarose were stiffened and largely unresponsive to quasi-static changes in alveolar pressure and excluded perfused fluorescent dye, in contrast to alveoli farther away from the blockage (**Fig. 5h**, **Supplemental Fig. 12, Supplemental Video 11**). This type of framework, where lung functionality can be locally arrested, could be a useful model of Hypoxic Pulmonary Vasoconstriction (HPV), where vessels contract in order to divert blood flow away from hypoxic alveoli^55,56^.

## Discussion

We have developed the crystal ribcage technology and demonstrated its capabilities for probing lung function and dysfunction across nearly the entire surface at the whole-organ, alveolar and cellular scales in real-time. The crystal ribcage enables multi-scale imaging of respiratory and circulatory functions while preserving the native 3-D architecture and cellular diversity of the lung in a wide array of pulmonary diseases (**Fig. 1**). In cancer, we found (i) differential remodeling of alveolar function in nodular vs infiltrative (co-optive) growth pattern, (ii) earliest stages of tumorigenesis that compromises alveolar respiratory function, and (iii) compromised capillary function around a nodular tumor (**Fig. 2**). We also demonstrated altered respiratory and circulatory functions in pneumonia, which affects delivery of oxygen, nutrient, and therapeutics, and impairs trafficking, recruitment, and clearance of immune cells (**Fig. 3**). By modulating vascular pressure in the lung, we found that neutrophils are strongly and reversibly mechano-responsive to vascular pressure in a mouse model of pneumonia and acute lung injury (**Fig. 4**). While more work needs to be done to understand the underlying mechanisms, the controllability of lung physiology in the crystal ribcage (e.g., vascular pressure in this application) provides enormous opportunity for understanding cause-and-effect in pulmonary pathogenesis. Lastly, we demonstrated that respiration-circulation coupling at the capillary level and found that this coupling is eliminated in acute injury (**Fig. 5**).

While the crystal ribcage supports probing of several lung diseases with parenchymal presentation, deeper imaging could expand the capabilities of our platform. Currently, any optical imaging modality is depth-limited in the lung due to light diffraction at the air-liquid interfaces of the alveoli^7,8^, a limitation unrelated to the crystal ribcage itself. When the alveoli are filled with liquid (e.g. edema) or cancer cells, light scattering is reduced which enables imaging up to 500 µm deep as opposed to 100 µm in air-filled alveoli (**Extended Data Fig. 6**). While the crystal ribcage allows dynamic imaging at physiological respiratory and circulatory rates (**Supplemental video 2,3**), we chose to probe the lung mechanics under quasi-static conditions to be able to report the accurate pressure values in the alveoli and capillaries. Ultrafast volumetric imaging techniques coupled with the crystal ribcage can resolve respiration and circulation dynamics at the “speed of life”, fast cell-cell communications such as calcium signaling, and quantify viscoelastic properties and energy dissipation at the alveolar level, all in a functioning lung. Additionally, recent optical imaging approaches^57^ to measure partial oxygen pressure could be leveraged to study gas transport in single alveoli^58^ in the crystal ribcage.

The crystal ribcage can also be used to image heart function in real-time. We imaged the pulsing of the heart atrium on the ventral surface of the crystal ribcage in response to elevated vascular pressure (**Extended Data Fig. 10, Supplemental Video 12**). Such experiments will allow further probing of circulation-respiration coupling in diseases such as pulmonary hypertension or arrhythmia in the future. Disease models, here, were developed *in vivo* on the order of days to weeks, and we probed the underlying physiology and pathology for 4-6 hours *ex vivo* inside the crystal ribcage, sufficient for studying dynamic events such respiratory function, immune cell migration, and circulation dynamics. However, recent studies have maintained human lungs up to 24 hours *ex vivo* ^59^, pig lungs up to 3 days^60^ and rats up to 7 days^61^. Utilizing these advances^62^, we will adapt the crystal ribcage to maintain lung viability for multi-day studies of disease dynamics. We are also taking advantage of the scalability of our fabrication process to develop the crystal ribcage for large animals such as pig and transplant-rejected human lungs^63^ to probe diseases and therapeutics in a more clinically relevant setting.

## Supporting information

Supplemental Video 1 | Functional ex vivo lung inside the crystal ribcage

Supplemental Video 2 | Whole lobe to single alveoli microscopy of lung under negative-pressure ventilation in the crystal ribcage

Supplemental Video 3 | Single alveoli microscopy of apex vs base in lung under negative-pressure ventilation in the crystal ribcage

Supplemental Video 4 | Crystal ribcage provides stable imaging surface for ventilated lung

Supplemental Video 5 | Immune cell migration in functional alveoli imaged in real-time through the crystal ribcage

Supplemental Video 6 | Apex to base image of lung with several metastatic tumors at alveolar resolution using OCT through the crystal ribcage

Supplemental Video 7 | Metastatic cancer cells remodel capillary function

Supplemental Video 8 | Real-time imaging of vascular transport at alveolar resolution in health and disease using crystal ribcage

Supplemental Video 9 | Neutrophil migration speed increases with higher vascular pressure

Supplemental Video 10 | Neutrophil extravasation and migration imaged through the crystal ribcage

Supplemental Video 11 | Local agarose blockage in the alveoli disrupts surrounding vascular dynamics

Supplemental Video 12 | Real-time imaging of spontaneous heart atrial pulsation in response to elevated pressure

Supplemental Video 13 | Air and perfusate leak from site of local excision in the lung

Supplemental Video 14 | Disrupted capillary flow in locally cauterized lung imaged through the crystal ribcage

## Acknowledgements

We thank the Neurophotonics Center at Boston University for their generous support and access to their facility. Adam Coats from the Hoffman lab at the University of Iowa provided the µCT image data sets used to fabricate the crystal ribcage. The research reported in this publication was supported by the Boston University Micro and Nano Imaging Facility and the Office of the Director, National Institutes of Health of the National Institutes of Health under award Number S10OD024993. The content is solely the responsibility of the authors and does not necessarily represent the official views of the National Institute of Health. Any opinion, findings, and conclusions or recommendations expressed in this material are those of the authors and do not necessarily reflect the views of the National Science Foundation.

## Author contributions

R.B., G.N.G. and H.T.N. conceived the project and wrote the manuscript; R.B. developed and validated the crystal ribcages and external devices suite, established transgenic mouse colonies, analyzed alveolar function, strain maps and collagen waviness; G.N.G. conducted the lung extraction and imaging, generated experimental mice, cultured cancer cells, and analyzed circulatory function data; L.S. analyzed perfusion, vascular transport, and neutrophil mechano-responsiveness data; D.S. analyzed alveolar and capillary function data, and maintained animal colony; R.L. developed registration and strain analysis algorithm; J.M. developed custom hardware for crystal ribcage and analyzed alveolar function data; C.E. developed the deformable crystal ribcage; B.E.H. generated the lobar pneumonia mouse model; J.L. and A.A.R. generated the fibrosis model; A.A.R. performed the histology and staining for fibrotic lungs; J.C.B. generated primary cancer mice, K.R. analyzed alveolar function data and cultured cancer cells; S. Zheng cultured cancer cells; S. Zhang fabricated fluorescent polyacrylamide beads; J.J. assisted in surgical techniques for *in situ* lung imaging; R.P. generated LPS induction model; K.T. provided transgenic MPR8-Cre^+^ mouse; S.M., G.L., J.P.M. and B.S. contributed to discussion on crucial aspects of the project; H.T.N. supervised the project and provided guidance on experimental design, data collection, data interpretation, and writing of the manuscript.

## Methods

### Mice

All experiments conformed to ethical principles and guidelines under protocols approved by the Boston University Institutional Animal Care and Use Committee. Mice were housed and bred under pathogen–free, ambient temperature, humidity, and light-dark conditions at the Boston University Animal Science Center, no housing or handling exceptions were made for this study. We used 6–12-week-old male and female mice for the experimental procedures. Female FVB mice aged 6-8 weeks (JAX #001800, Jackson Labs, ME) were purchased. A breeding pair of transgenic B6.129(Cg)-Gt(ROSA)26Sortm4(ACTB-tdTomato,-EGFP)Luo/J (JAX #007676, Jackson lab, ME)^64^, hereafter referred to as mTmG, was purchased to start a colony and was the primary source of all animals for the experiments. Male neutrophil reporter mice B6.Cg-Tg(S100A8-cre,-EGFP)1Ilw/J (JAX #021614)^65^ were purchased from JAX and crossed with in-house homozygous mTmG females to breed MRP8-mTmG mice that were used for neutrophil migration dynamics. We used a 6-week SPC-GFP mouse (Jax #028356) to visualize alveolar epithelial type II cells.

### Metastatic cancer model

E0771-H2B-dendra2 mouse cancer cells or MCa-M3C-H2B-dendra2 (both gifts from Rakesh Jain) were used to model breast cancer metastasis to the lung in mTmG and FVB mice, respectively. Cells were cultured in DMEM (Corning) plus 10% fetal bovine serum (Gibco) plus 1% penicillin/streptomycin (Gibco). Cells were harvested at ∼80% confluency, washed twice with phosphate buffered saline (PBS), counted, and resuspended in DMEM prior to tail vein injection into mice at a concentration of 2 million per 300uL complete medium. All cell lines tested negative for mycoplasma (Mycoalert Plus Mycoplasma Detection Kit, Lonza, Allendale, NJ), and authenticated before use by IDEXX laboratories (North Grafton, MA). Mice injected with E0771-H2B-dendra2 were sacrificed on days 5-7 post injection for microscopy of mature nodules. Mice injected with MCa-M3C-H2B-dendra2 were sacrificed on days 10-14 for microscopy of mature nodules. Largest tumor nodules studied were 2 mm in diameter and mice were excluded from the study if they presented with labored breathing, hunched posture, or ruffled fur due to tumor progression (as stipulated by BU IACUC). For single cell deformation studies, E0771-H2B-dendra2 cells were plasma membrane labeled with 1,1’-Dioctadecyl-3,3,3’,3’-Tetramethylindocarbocyanine, 4-Chlorobenzene Sulfonate Salt (DiD, Invitrogen) before imaging, then washed twice with complete cell culture medium and injected intracardiac into the mouse at a concentration of 6 million per 300uL solution. To ensure that single cells were sequestered inside of blood vessels for single cell deformation studies, we tail vein injected a 50 µL volume of bovine FITC-albumin (5 mg/ml, ThermoFisher Scientific, A23015) approximately 10 minutes before intracardiac injection of single cells to label the blood vessel lumens.

### Bacterial pneumonia model

We induced bacterial pneumonia in mice as previously described^66^. Briefly, *S. pneumoniae* serotype 3 (Sp3) (ATCC; Manassas, VA) was cultured on plates of BBL Trypticase Soy Agar with 5% sheep blood (BD Trypticase Soy Agar II) (BD Biosciences) in a 37°C humidified 5% CO2 environment for 10-11 hours and diluted in 0.9% sodium chloride to the desired optical density. Mice were anesthetized using a ketamine/xylazine cocktail (100 and 10 mg/kg, respectively) and approximately two million CFU in 50 μL were instilled into the left lung lobe via the left bronchus, accessed through the surgically exposed trachea. Mice were sacrificed 24-36 hours post induction to image the acutely injured lung.

### Pulmonary edema visualization

Evans blue has historically been used as a label for vascular leakage and pulmonary edema in rodent models of pulmonary disease as it binds serum albumin^67-69^. We injected 50 µl of 10 mg/ml solution of Evans blue (ThermoFisher Scientific) with 100 µL of 1.25 mg/ml heparin saline into the tail vein of mice 3-4 hours prior to excision of Sp3 infected lungs. Even after pulmonary perfusion with unlabeled RPMI media, edematous regions of the lung retained the characteristic blue color and far-red fluorescence of Evans blue (Fig. 1e, Extended Data Fig. 9, Supplemental Fig. 8)

### Primary cancer model

We induced tumorigenesis in 6-week female A/J mice as described previously^70^. Mice were treated with a cocktail of Benzo[a]pyrene (BaP) (B1760, Millipore-Sigma, USA) (0.5 mg/dose) and 4-(Methylnitrosoamino)-1-(3-pyridinyl)-1-butanone (NNK) (78013, Millipore-Sigma) (0.41 mg/dose) were dissolved in cotton oil (C7767, Millipore-Sigma) and delivered to the back of the tongue using a metal 20G feeding tube, twice per week for 4 weeks and sacrificed at 20 weeks to image the lung in the crystal ribcage.

### LPS acute inflammation model

mTmG mice were treated with 100 µg *Escherichia coli* (*E. coli*, serotype O55:B5, Sigma-Aldrich) lipopolysaccharide (LPS) diluted to 100 ug/60 µL in PBS via intratracheal injection using an angio-catheter as previously described^71^, to model acute injury. Mice were sacrificed 3-4 hours post induction for microscopy of the acutely injured lung.

### Pulmonary fibrosis model

Bleomycin was delivered to the lungs to generate a model of pulmonary fibrosis as previously described^72^. Mice were anesthetized with ketamine/xylazine cocktail were given 1 U/Kg bleomycin (APP Pharmaceutical, LCC Schaumburg, IL, USA) intratracheally using a 30G insulin syringe. Mice were sacrificed 13-14 days after injection to image fibrotic regions. Collagen was imaged using 2-photon second harmonic generated signal (SHG) and validated by Masson’s trichrome, and Col1A1 (E8F4L) XP Rabbit mAb cell signaling (Catalog 72026, ThermoFisher Scientific) with Donkey anti-Rabbit IgG (H+L) highly cross-adsorbed secondary antibody alexa fluor 555 (Catalog A-31572, ThermoFisher Scientific) staining of the same spatial locations (**Supplemental Fig. 3)**

### Elastase induced emphysema model

COPD was induced in mTmG mice by digesting the elastase in the lung using porcine pancreatic elastase (PPE) (EC134 Elastin Product Company Inc., Owensville, MO). A previously published protocol was followed for single dose treatment^73^, briefly 6 IU PPE was dissolved in 100 μL saline and delivered to the mouse (23-25 g). The solution was instilled at the back of the tongue of anesthetized and vertically suspended mice, after which it was allowed to recover normally. Mice were sacrificed at 21 days from instillation to image the lung inside the crystal ribcage.

### Histological staining for fibrosis evaluation

Lungs were perfused with 10 mL of sterile cold PBS to remove blood then inflated with 4% Paraformaldehyde (PFA). The trachea was clogged with a thread and lungs were fixed in 4% PFA for 24 h at 24 °C. Fixed lungs were washed in 1x PBS three times 5 min at each interval at 24 °C. Fixed lungs were dehydrated in 50%, 70%, 85%, 95% and 2x 100% ethanol, cleared in 3x Histo-Clear (a xylene substitute, National Diagnostics) and embedded in paraffin. Formalin-fixed paraffin-embedded (FFPE) tissues were cut to serial sections (5 μm). FFPE sections were deparaffinized using a standard protocol of Histo-Clear and alcohol gradients. Masson’s trichrome staining was performed by using a commercially available kit (HT15-1KT, Sigma-Aldrich). Hematoxylin and Eosin (H&E) staining was performed using Harris Hematoxylin solution (ThermoFisher Scientific) and Eosin-Y solution (ThermoFisher Scientific). After dehydration in 70%, 95%, and 100% ethanol series and clearance with Histo-Clear, the stained slides were mounted and imaged using Olympus BH-2 light microscope (Olympus).

### Immunofluorescence staining for fibrosis samples

FFPE sections were deparaffinized using a standard protocol of Histo-Clear and alcohol gradients. Antigen retrieval was performed by boiling the slides in citrate buffer (10 mM citric acid, 0.05% Tween-20, pH 6.0) for 20 minutes, and then, the slides were allowed to cool at 24 °C for at least 30 min. The slides were washed with PBS and permeabilized with 0.2% Triton-X for 15 minutes. Following blocking in 5% normal donkey serum in PBS containing 0.2% Triton-X for 2 hours at 24 °C. The slides were incubated with an antibody against COL1A1 (E8F4L rabbit mAb, 1:200, Cell signaling Technology, #72026T) followed by a fluorescence-conjugated secondary antibody (Donkey anti-Rabbit IgG (H+L) Highly Cross-adsorbed Secondary Antibody, Alexa Fluor 555, ThermoFisher Scientific #A-31572, MA, USA, 1:1000 dilution) and DAPI (ThermoFisher Scientific #62248, Waltham, MA, USA, 1:1000 dilution) to counterstain nuclei. Stained slides were mounted with Aqua-Poly/Mount (Polysciences, Warrington, PA, USA) and imaged using Olympus CKX53 inverted microscope (Olympus).

### *In situ* lung imaging

Mice were anesthetized as described above and ventilated through a tracheal cannula to prevent lung collapse before sacrificing the mouse. The skin and fascia were dissected the intercostal muscles between the ribs was carefully separated under a stereomicroscope to expose the ribcage pleura. A 2mm x 3mm retaining clamp was inserted in the cleared section to prevent the leftover tissue strands blocking the opening. The PEEP on the lung was altered to confirm that the lung-maintained contact with the internal pleural membrane and that the lung was responsive to pressure changes. The mouse chest carcass was angled upwards for imaging.

### Lung preparation

Isolated mouse lungs were ventilated (Kent Physiosuite Mouse ventilator, Kent Scientific) through a tracheal cannula and perfused through two cannulas inserted into the pulmonary artery (PA) and left atrium as previously described^44^. Perfusates included whole blood, cellular, or acellular media containing fluorescent labels for vascular lumens and circulating cells. After cannulation, the lung-heart bloc was placed into the crystal ribcage for microscopy under controlled ventilation/circulation parameters. Perfusion pressure was controlled using a 60 mL reservoir containing the perfusate medium (serum-free RPMI, Gibco) suspended at different hydrostatic heights above the lung.

### Lung microscopy

Prepared ex vivo lungs inside the crystal ribcage were imaged under quasi-static inflation conditions or during dynamic ventilation using an inverted laser-scanning confocal microscope (Olympus FV 3000) under objective magnifications of 1.25x, 10x, and 60x, with environmental temperature control set to 37℃ using FluoView (v2.5.1.228) acquisition software. The confocal microscope was used with a resonant scanning mirror to increase the imaging speed up to 15 framer per second (FPS) for imaging the dynamically ventilated lung, which increases speed by compromising some of the image resolution. Additionally, lungs were imaged using an upright Nikon fluorescent stereomicroscope, upright Nikon CSU X1 spinning disk confocal with 1x, 2x, 4x and 10x objectives, ThorLabs upright optical coherence tomography probe with a 10x objective and an upright Bruker two-photon microscope (using Prairie View acquisition software) equipped with a 16x water immersion objective. Microscopy data was visualized and processed using FIJI and MATLAB R2021b, R2022a, R2022b.

### Crystal ribcage development and fabrication

The entire development of the crystal ribcage is described as a supplemental protocol^74^. Briefly, previously recorded µCT scans of C57B/6 and AJ mice chest cavities (courtesy the Hoffman group at the University of Iowa^75-77^), and FVB mouse chest scans (performed at the Boston University Micro-CT imaging facility) were segmented in MATLAB (R2019b Mathworks, Natick MA) and imported into Solidworks 2019 (Dassault systems, France) to create a refined geometry of the chest.The model was made age-specific by scaling the binary object by mouse lung volume reported by age^78^ (**Extended Data Fig. 1**) and printed in FormLabs (Somerville, MA) clear resin on a Form3 printer. The mold was polished to get a high gloss surface finished buck. A sacrificial sugar mold was cast from the buck to make semi-flexible polymethyldisiloxane (PDMS) ribcage. The buck was used directly to thermoform polystyrene sheets over it to make the rigid crystal ribcage. The internal surface of the PDMS crystal ribcage was treated with a 1% PDMS-PEG Block Copolymer (DBE-712, Gelest Inc., Morrisville, PA) and allowed to cure to improve hydrophilicity as previously described ^79^.The rigid crystal ribcage was treated in an oxygen plasma cleaner to make the surface hydrophilic.

### Quasi-static alveolar diameter measurement

Microscopy images recorded with a 10x objective and an additional 1.5x optical zoom were loaded into FIJI-ImageJ and sum-wise projected in the z-direction for all pressure conditions. Alveoli common to all quasi-static or dynamic pressures were annotated prior to making measurements. The mean alveolar diameter was calculated by recording 2 length values between anatomical features that were diametrically opposite to each other. The mean lengths were recorded for all corresponding alveoli at different pressures. These measurements were compiled in MATLAB R2021b to make raincloud plots^80^ and compute group-wise statistics.

### Alveolar septum strain calculation

We used a previously published deformable registration algorithm, deedsBCV^81^ to autonomously determine the displacement map between images of the lung tissue at different quasi-static inflation pressures, here alveolar pressures of 7 and 12 cmH_2_O. We subsequently used a custom algorithm (finite difference methods are subject to noise that obscures signal) to compute the strain map (Supplemental Fig. 6) with a cubic smoothing spline to approximate the displacement map and evaluate its gradient derivative and calculate the canonical strain tensor components directly. This method gives us a displacement gradient continuous in its second derivatives which avoids obscuring the true signal in the noise. We validated the obtained displacement maps against known displacements applied to images being registered. We validated the effect of the smoothing spline with multiple kernel sizes to ensure that the final registered image matches the target image. The final strain computation algorithm was validated for known synthetic displacement maps to ensure parity between the mathematical ground truth and image-based strain estimation.

### Dynamic alveolar diameter measurement

The 2-D image series of the lung were loaded in MATLAB R2022a and segmented by applying a global threshold to each image. The binarized images were analyzed using the built-in *regionprops* function to obtain the equivalent diameter of a circle with the same area as each binarized segment. Diameters greater than 500 μm or less than 10 μm were ignored. The mean diameter of each image was plotted as a time series and the peaks of the curve were obtained to find when the cycle was repeated. The total number of frames were linearly distributed to obtain a normalized ventilation cycle and pooled to get a single mean alveolar diameter curve per ROI to obtain the width at half-maximum of the ventilation cycle for each ventilation rate.

### Collagen waviness quantification

The 3-D images of the tumor nodules were imaged with a Bruker 2-photon microscope with the laser excitation set at 920 nm. The dendra signal from cancer cells was collected through a 525/50 nm emission filter and the second harmonic generation collagen signal was collected through a 460/50 nm emission filter on separate channels. The speckle noise was removed from the 3-D stack using a 2-pixel median filter in MATLAB R2021b medfilt2() and manual traces were made along the collagen fibers between successive nodes. The total path length traced was divided by the end-end distance of the start and end points of the trace to determine the waviness index.

### Single cell measurement

Microscopy images at each quasi-static pressure condition were loaded into FIJI-ImageJ and cells were traced around manually. Circularity of single cancer cells was calculated as 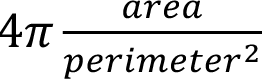 The tabulated results were compiled in MATLAB R2021b to perform statistical comparisons between the groups of cells at each condition. We confirmed that single cells were sequestered inside blood vessels by exclusion of FITC-albumin dye, which was injected into the mouse via the tail vein (50 µL of 10 mg/ml FITC-albumin in saline) prior to injection of singularized cancer cells.

### Cancer nodule size estimation

We estimated the size of cancer nodules for the analysis reported in Fig. 2h by first assuming an average cell size of 10 µm and measuring the diameter of the cancer nodule in in the XY plane, from confocal microscopy images of nodule-bearing lungs in the crystal ribcage from confocal microscopy images (representative image in Fig. 2a). We estimated the size of the nodules in terms of cell number by dividing the diameter of the nodule by average cell size.

### Perfusion and vascular transport data analysis

We extracted the mean fluorescent intensity across all pixels in the perfusion timelapse video, at each time frame for each ROI. Then, we normalized the mean fluorescent intensity curve of each ROI to the maximum of the mean intensity from all ROIs. Multiple replicates for each ROI are combined to produce a final normalized mean intensity (solid line) and the respective standard error of the mean (shaded areas). The duration of dye intensity greater than 50% of the normalized maximum intensity was reported as T_50_. The time from initial increase in pixel intensity time series to the normalized maximum intensity was calculated (T_max_). Step bolus injections for tumor diffusion were processed in ImageJ to obtain dye intensity intratumor and normalized to baseline intensity measured within the vasculature.

### Cell tracking for neutrophil migration

Post-processing of neutrophil migration data was performed in MATLAB2021b. Multi-channel recordings from microscopy were stabilized and single neutrophil centroids were computed per frame to obtain corresponding position and speed information. Persistence was calculated as the total displacement divided by contour length of the path. Where relevant, migration through the intravascular space was defined as movement confined within the vasculature, intra-alveolar as movement within alveolar spaces, and interstitial as those crossing between vascular and alveolar spaces. Neutrophil shape was segmented using an intensity threshold and the *regionprops* function was used to compute aspect ratio and circularity.

### Statistics and Reproducibility

No statistical method was used to predetermine sample size and no data were excluded from the analyses. The experiments were not randomized, and investigators were not blinded to allocation during experiments and outcome assessment. All experiments were performed at least twice in separate mice to generate biologically independent samples. Data are presented as mean ± standard error of the mean (SEM). All statistics presented here were calculated as a two-tailed Student’s t-test to determine significance or p-values between groups of data. All p-values are reported on figures for the users to observe statistical trends of data. Significant differences were considered when p < 0.05. Statistical analyses were performed in MATLAB R2021b and R2022b.

**Data Availability:** A demonstration data set has been deposited in a Zenodo repository^82^ and are available from DOI: doi.org/10.5281/zenodo.7939072. Additional microscopy data is available on request from the corresponding author.

**Code availability:** The source code and algorithms used in this study have been deposited in a Zenodo repository^82^ and are available from DOI: doi.org/10.5281/zenodo.7939072 licensed under the CC BY 4.0 for use in research.

**Extended Data Figure 1.**
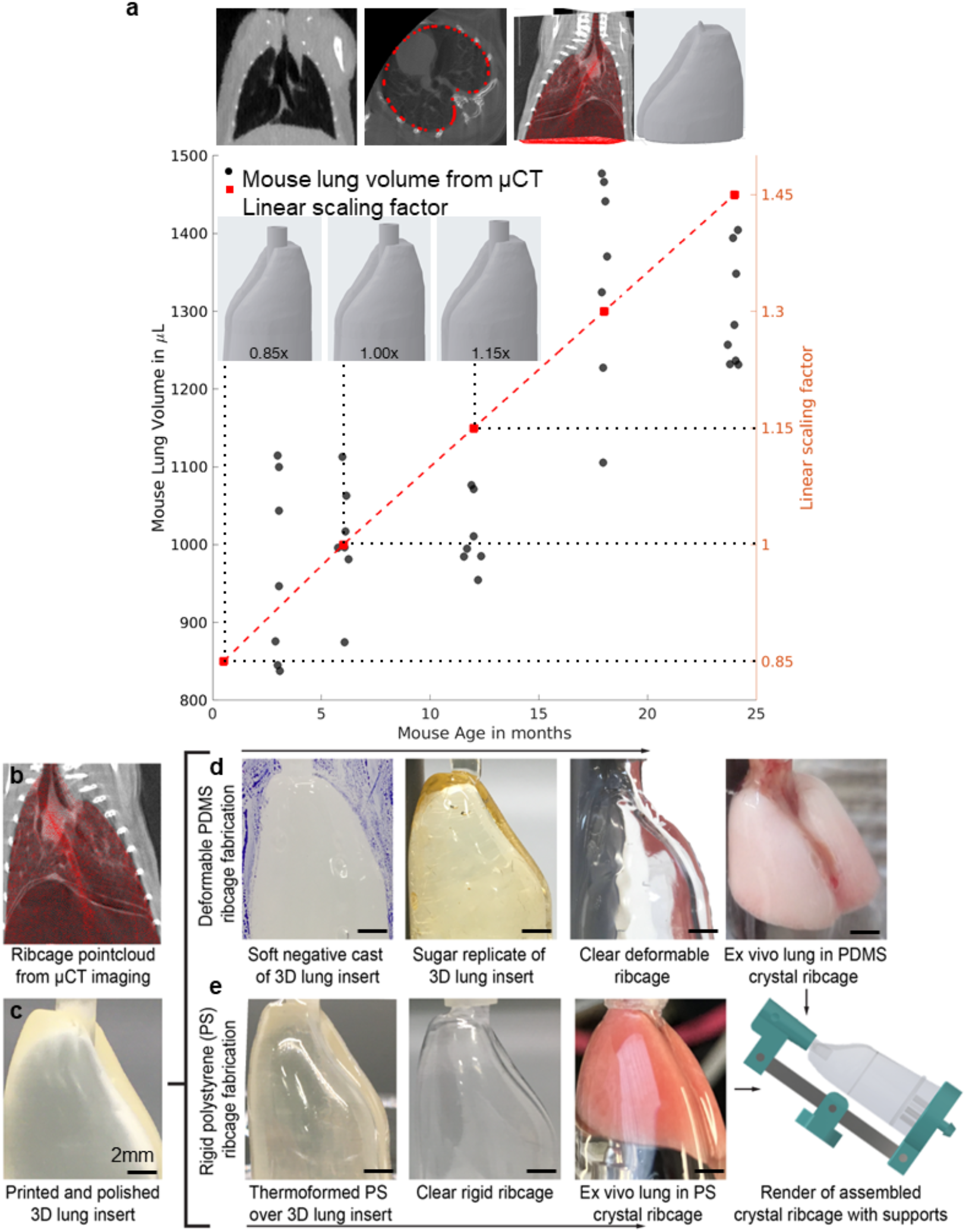
| Mouse strain specific µCT informed 3D molds scaled by age and fabrication of crystal ribcages. **(a)** Recreating mouse ribcage geometry from µCT data^75-77^, here shown for C57/B6 mice. Top: Sagittal view of mouse chest with fully inflated lung (dark region), distribution of axial control points to recapitulate ribcage cross-section and comparison of reconstructed geometry overlaid on sagittal and coronal µCT slices. Lung volume variation^78^ with age was normalized against the mean volume at 3 months to construct a linear fit for scaling 3D lung inserts by age. (**b-e**) 3D printed models were used to make both PDMS and polystyrene crystal ribcages in two independent processes^74^.

**Extended Data Figure 2.**
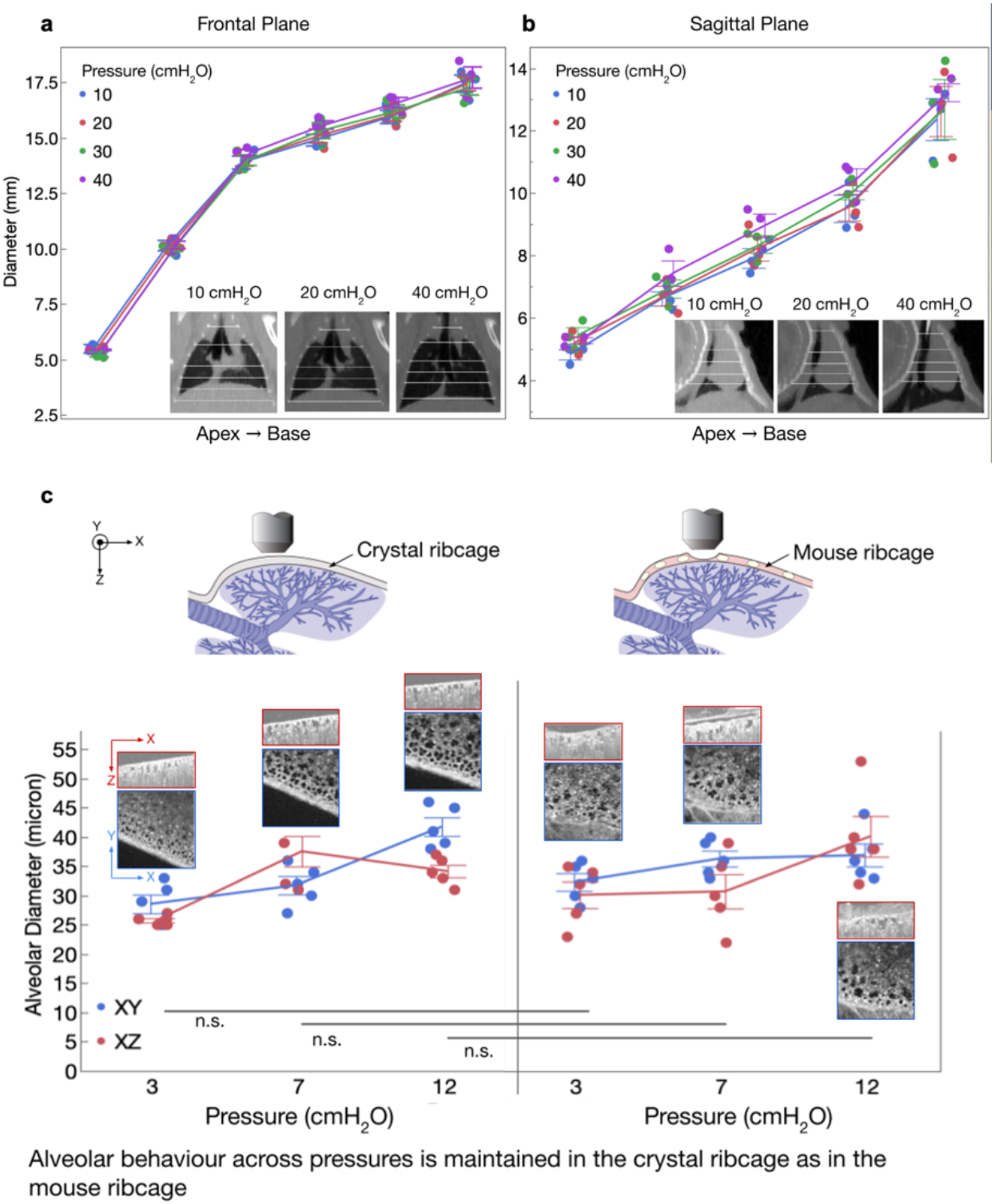
| Native mouse ribcage does not show significant variation in diameter across its anatomy and alveolar function (diameter in μm) remains unchanged between crystal ribcage and mouse ribcage. Previous µCT data^75-77^ used to define the mouse chest cavity was also used to (**a**) quantify the expansion of the native mouse ribcage as a function of pressure and anatomical location during *in situ* ventilation. There was no significant difference between the ribcage diameter at tracheal pressures ranging from 10 to 40 cmH_2_O in both the frontal and sagittal planes from the apex to the base of the ribcage. The increase in volume with increasing pressure was accommodated by the increasing length of the lung (movement of diaphragm and abdomen) and not the width of the ribcage. This confirms that there was little contribution of the changing ribcage diameter to lung volume change during normal physiological conditions *in situ*. The scatter of the data over n=3 mice is presented along with the mean ± SEM traces. (**b**) A mouse ribcage was thinned down to make a window above the pleural sheath to image the functional lung using OCT, the same lung was then measured in the crystal ribcage for the same changes in pressure. There was no significant difference between alveoli function (measured as changing diameter) across the pressure range for m=5 ROI over n=1 mouse. The scatter of the data is presented along with the mean ± SEM traces. Comparison between groups performed with two-tailed Student’s t-test for significance < 0.05.

**Extended Data Figure 3.**
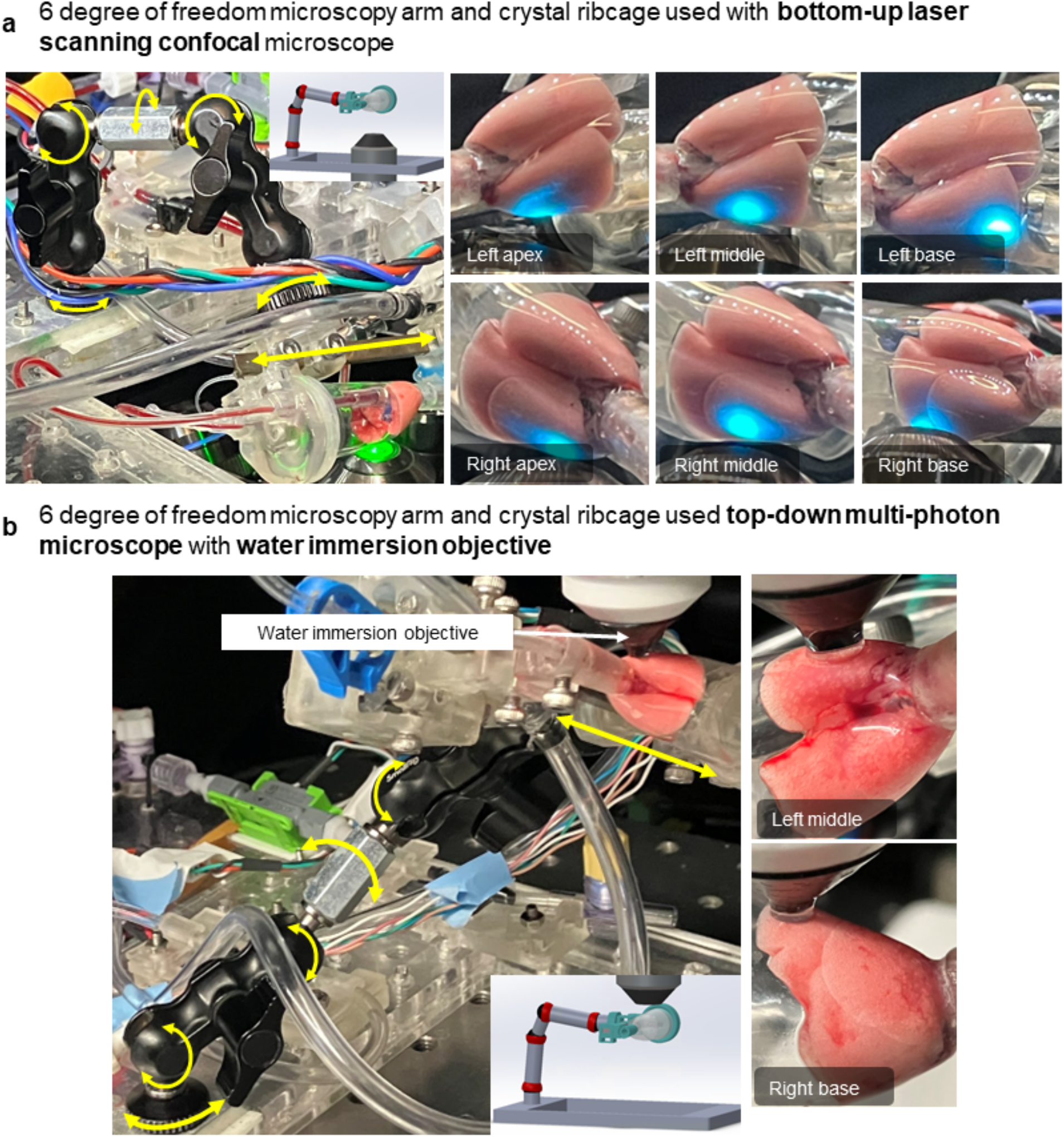
| Experimental setup and 3D renders of the microscopy arms used to image different orientations of the crystal ribcage with upright and inverted microscope configurations. We successfully used the stage to image (**a**) with an inverted Olympus Fluoview 3000 laser scanning confocal microscope and (**b**) with upright microscopes, such as the ThorLabs optical coherence tomography probe and Bruker multi-photon microscope. The crystal ribcage is microscope agnostic and compatible with both air and water-immersion objective in both configurations.

**Extended Data Figure 4.**
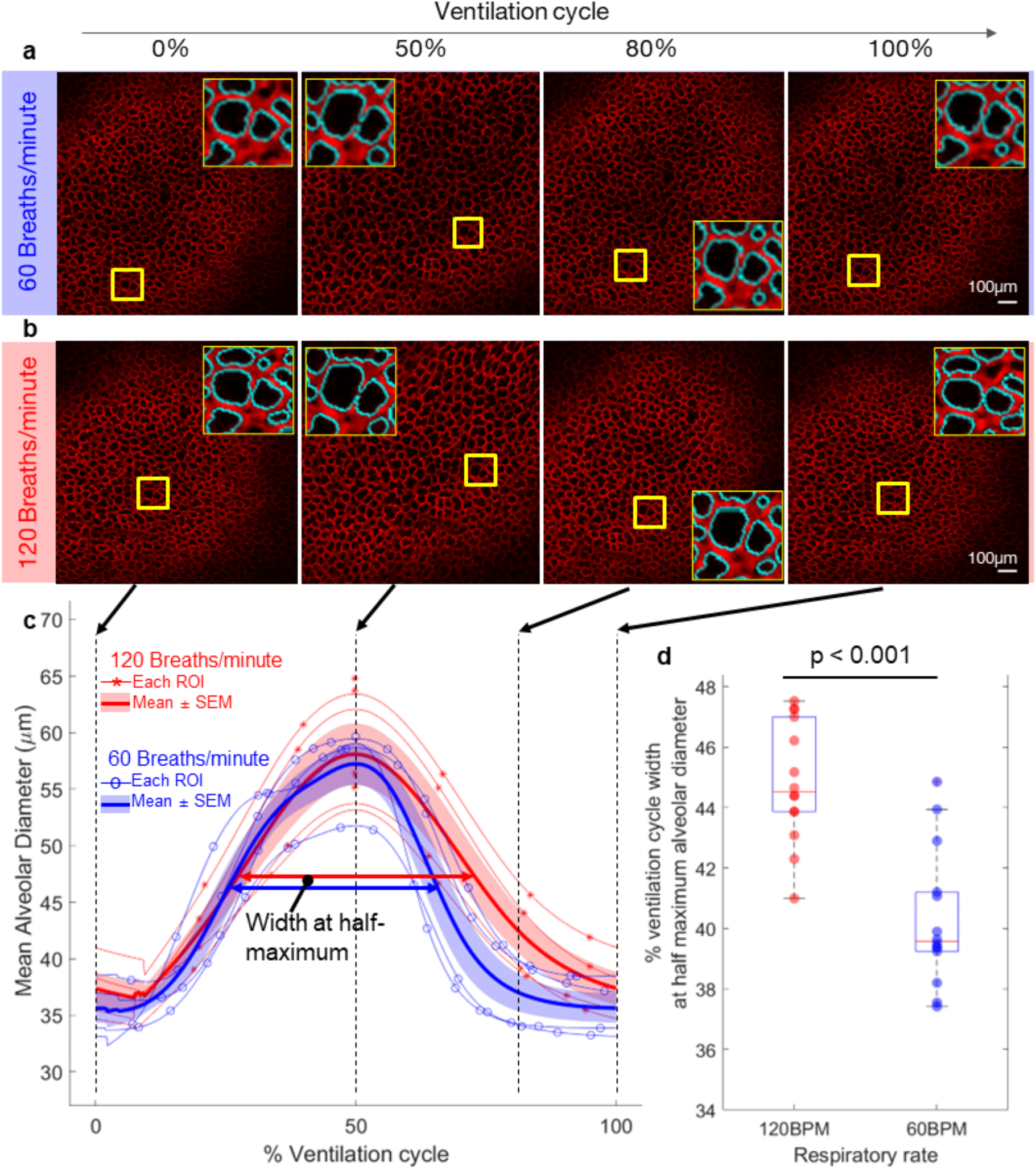
| Dynamic ventilation of mouse lung shows alveoli remain at a larger diameter for longer duration at higher ventilation rate. Comparing the same region of the lungs ventilated at (**a**) 60 breaths/minute with (**b**) 120 breaths/minute by (**c**) the mean ± SEM alveolar diameter from m=4 ROI over n=2 mice with a normalized ventilation cycle. Blue traces in the inset images show the segmentation boundary drawn to quantify single alveoli. (**d**) The width of the normalized ventilation cycle from m=4 ROI over n=2 mice at each respiratory rate are statistically larger at the higher respiratory rate. Boxplots present median with 25^th^ and 75^th^ percentiles, whiskers are the maximum and minimum data points not considered outliers. Comparison between groups performed with two-tailed Student’s t-test for significance < 0.05.

**Extended Data Figure 5.**
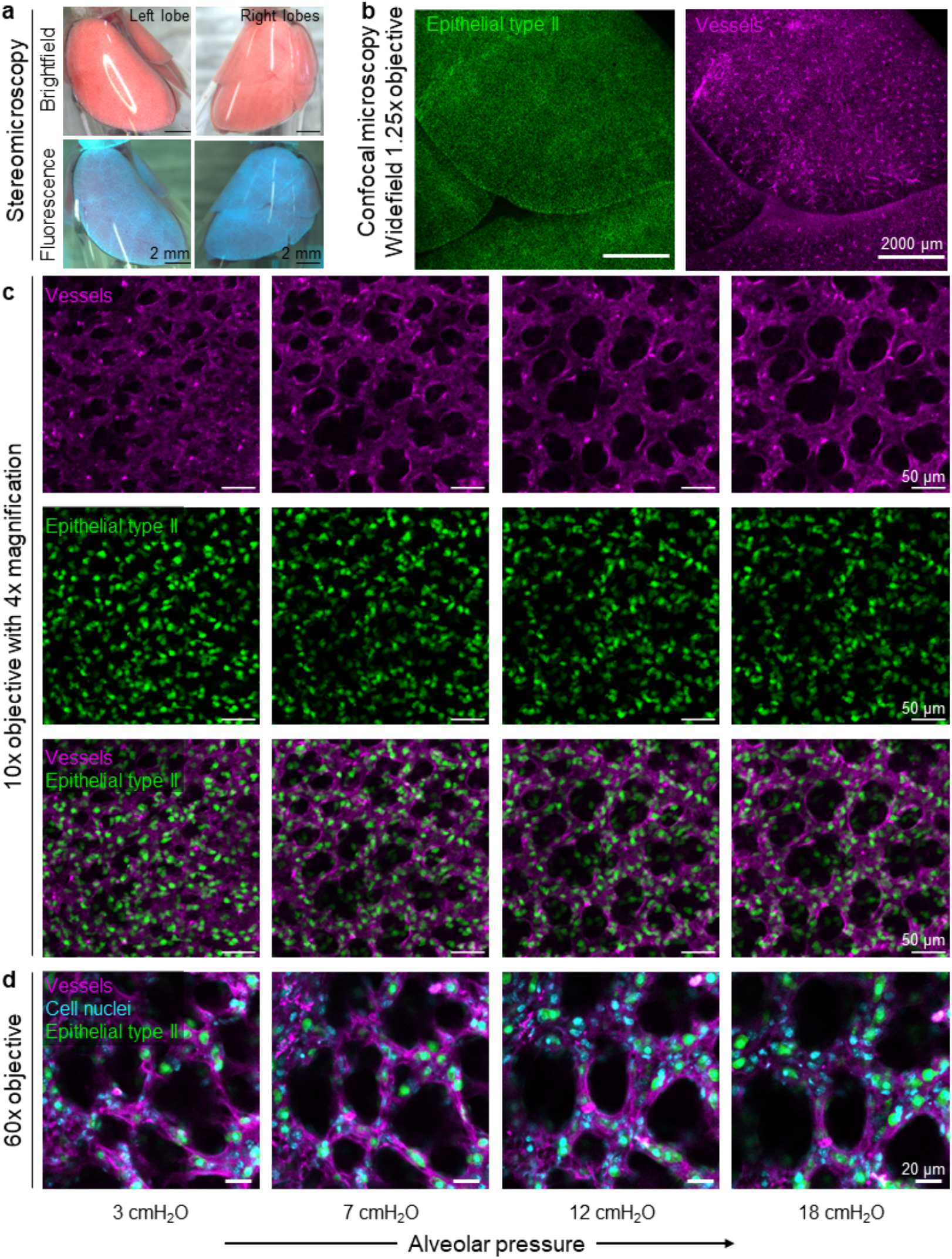
| Resident alveolar type II cells, with GFP-labelled surfactant protein C (green) can be imaged inside the crystal ribcage. (**a**) Stereomicroscope fitted with a NightSea GFP filter to visualize the SPC-GFP distribution at the whole organ scale. (**b**) Confocal microscopy of the SPC-GFP population at the lobe scale on the right side of the lung at 1.25x magnification. Vessels are labeled by injection of 50 µL of 10 mg/ml Evans blue dye in saline *in vivo* prior to extraction of the lungs. (**c**) Confocal microscopy at the 10x magnification shows single SPC-GFP+ cells in the lung distal parenchyma. (**d**) Single cell nuclei and SPC-GFP signal in cell cytoplasm visible at single cell resolution using a 60x water immersion lens.

**Extended Data Figure 6.**
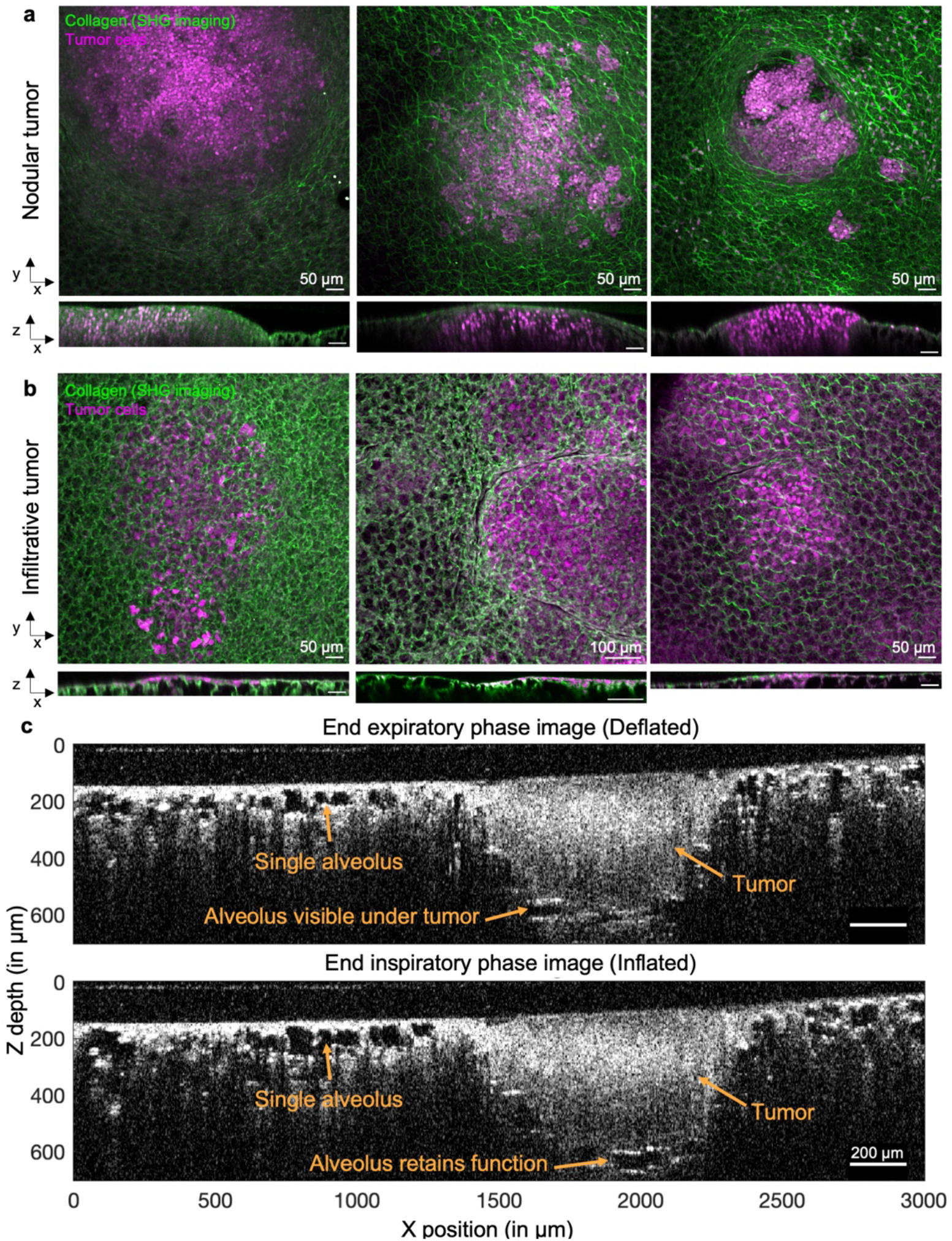
| Orthogonal views comparing the spatial distribution of nodular and infiltrative tumors, for three separate instances of each phenotype and solid phase nodular growth tumor increases imaging depth in sagittal image section using optical coherence tomography. (**a**) Nodular tumors extended into the lung tissue to greater depths and bulge out of the pleural surface. (**b**) Infiltrative tumors are located more superficially on the pleural surface. (**c**) Alveolar function deeper than 100µm from the surface can be imaged through the tumor nodule up to a depth of 400µm from the surface, as the gas phase of the alveoli is replaced by the solid phase tumor, which reduces light scattering.

**Extended Data Figure 7.**
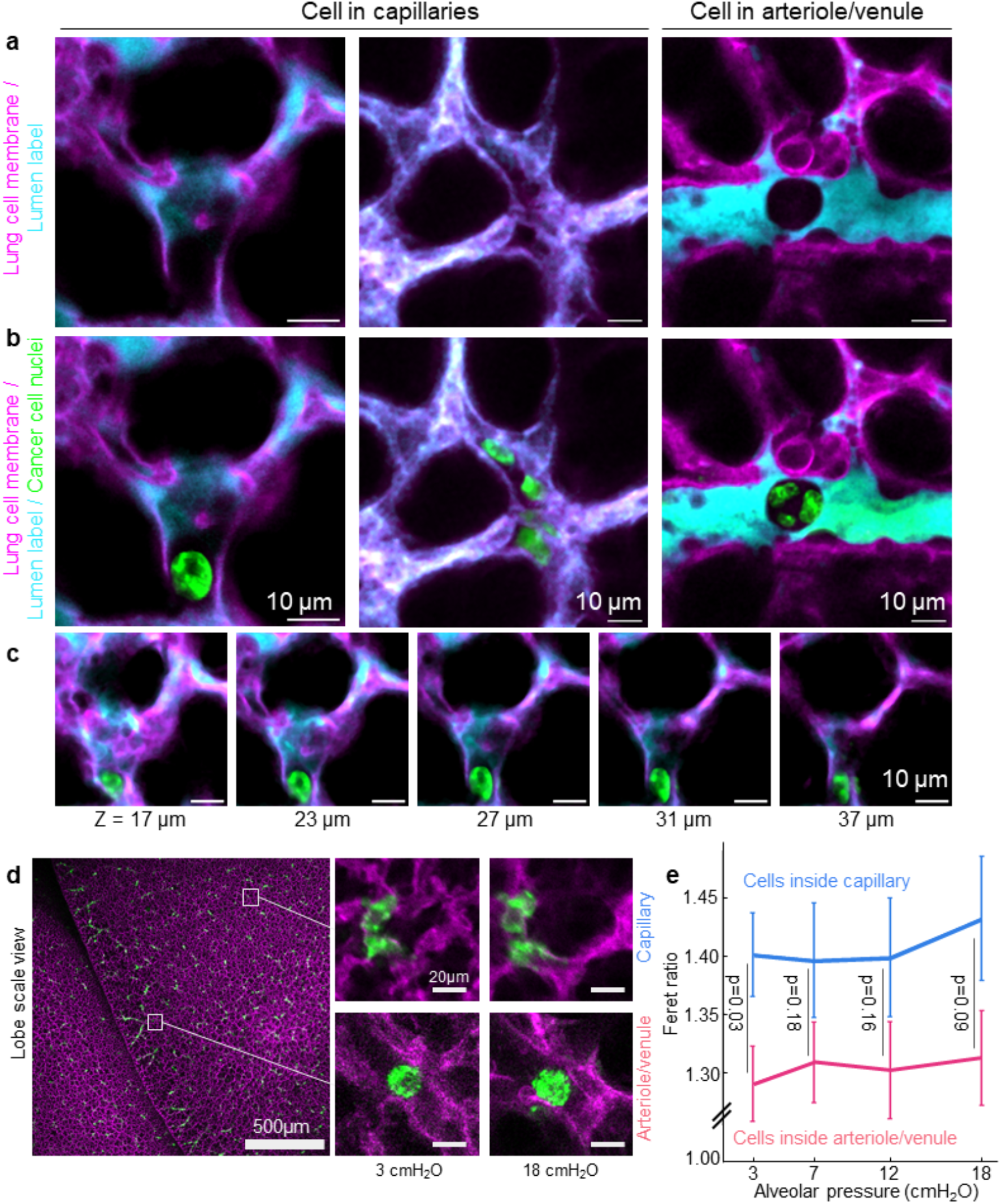
| Single cell deformation in response to changing alveolar pressure. (**a**) Cell-scale confocal microscopy of the lung inside the crystal ribcage with a FITC-albumin vascular lumen label showed the shadow of the cancer cell (**b**) which was imaged in a separate channel and merged to show cancer cells stretched in capillaries compared to arterioles and venules. (**c**) XY-views of the cancer cell at different depths of confocal imaging to show the cell is completely enclosed in the capillary. (**d, e**) Cells in capillaries quantified by the Feret ratio trended towards greater deformation in comparison to cells in larger arterioles-venules. Feret ratio quantified for n=40 cells in capillaries, n=30 cells in arteriole-venule at each alveolar pressure and data presented as mean ± S.E.M.

**Extended Data Figure 8.**
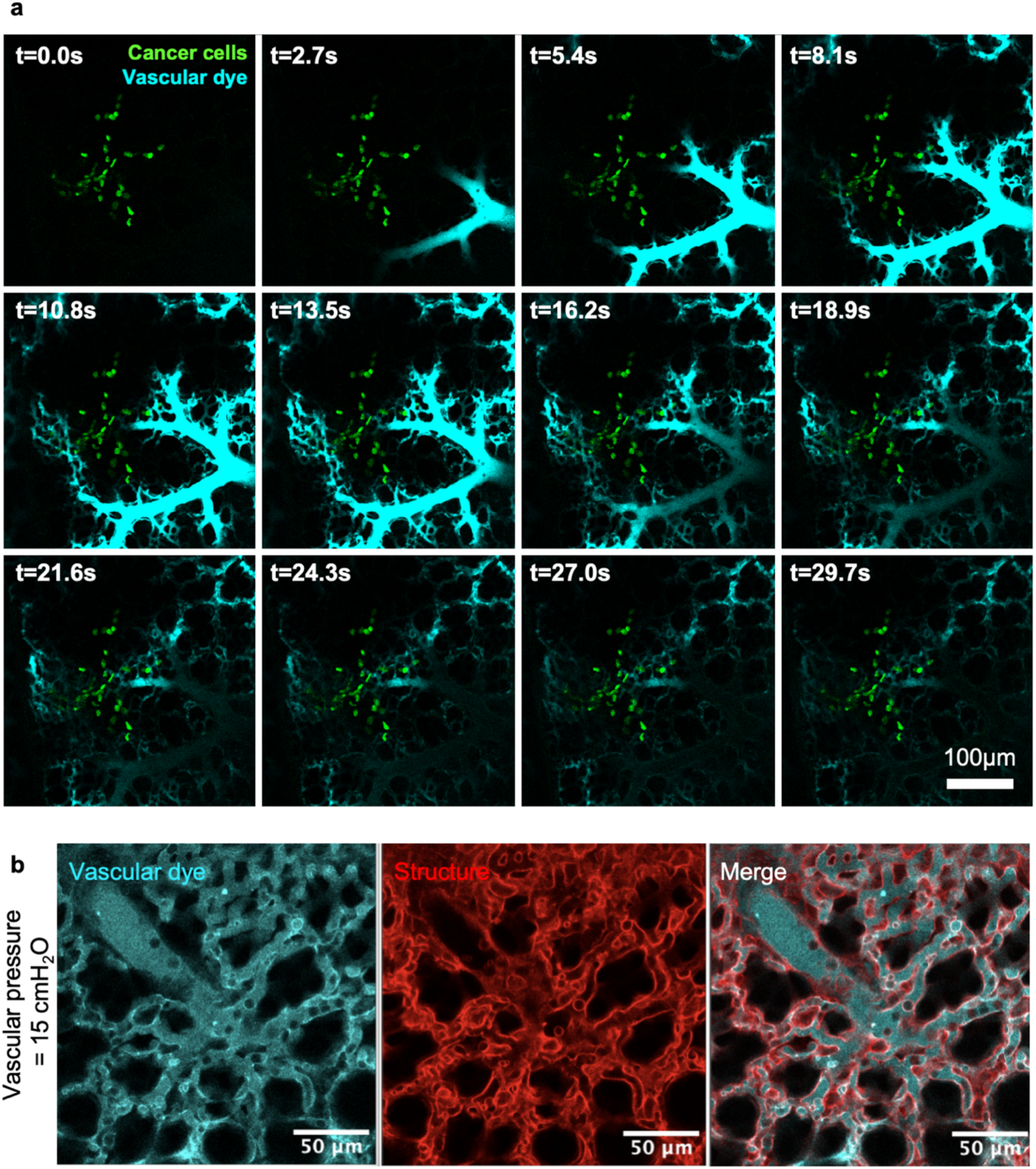
| Metastatic cancer cells disrupt vascular dye distribution in capillaries and the crystal ribcage used to visualize individual lung capillary walls and perfused lumens at high magnification (60x). (**a**) Capillary scale confocal microscopy imaging of single cancer cells that disrupt the distribution of dye observed over a time lapse imaging. (**b**) Vascular lumens were labeled with 10 mg/ml FITC-albumin dissolved in saline, injected intravenous prior to excision of the lungs for high-resolution confocal microscopy with a 60x objective.

**Extended Data Figure 9.**
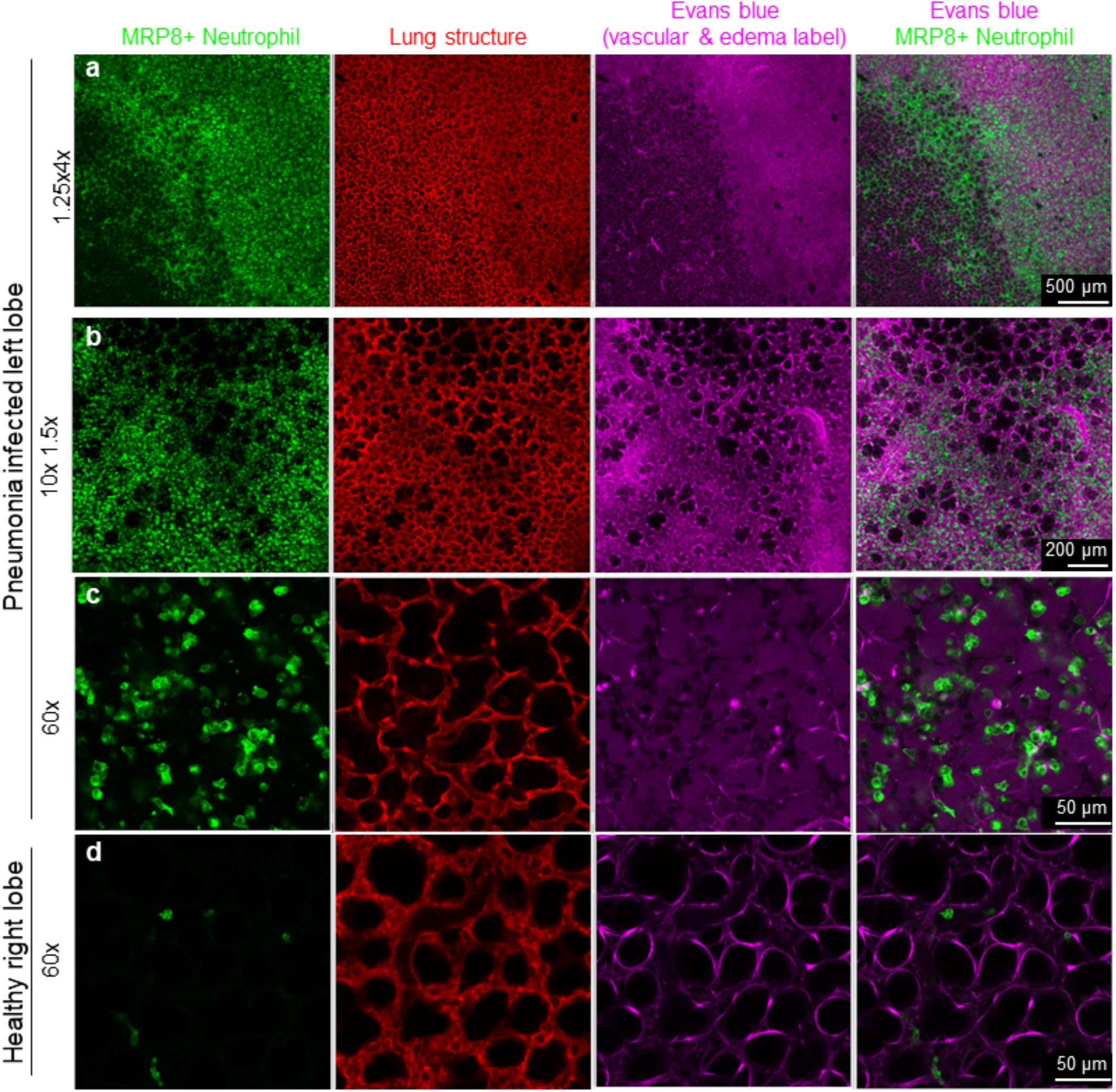
| MRP8+ Neutrophil cells co-localized with edema in lungs after at 30 hours lobar pneumonia injury. Correlation of the edema label with neutrophil cells is imaged (**a**) using 1.25x, (**b**) 10x and (**c**) 60x objectives to confirm that all alveoli with neutrophils (green) are edematous (magenta). (**d**) The contralateral lobe was also imaged to show the absence of edema and neutrophils inside alveoli.

**Extended Data Figure 10.**
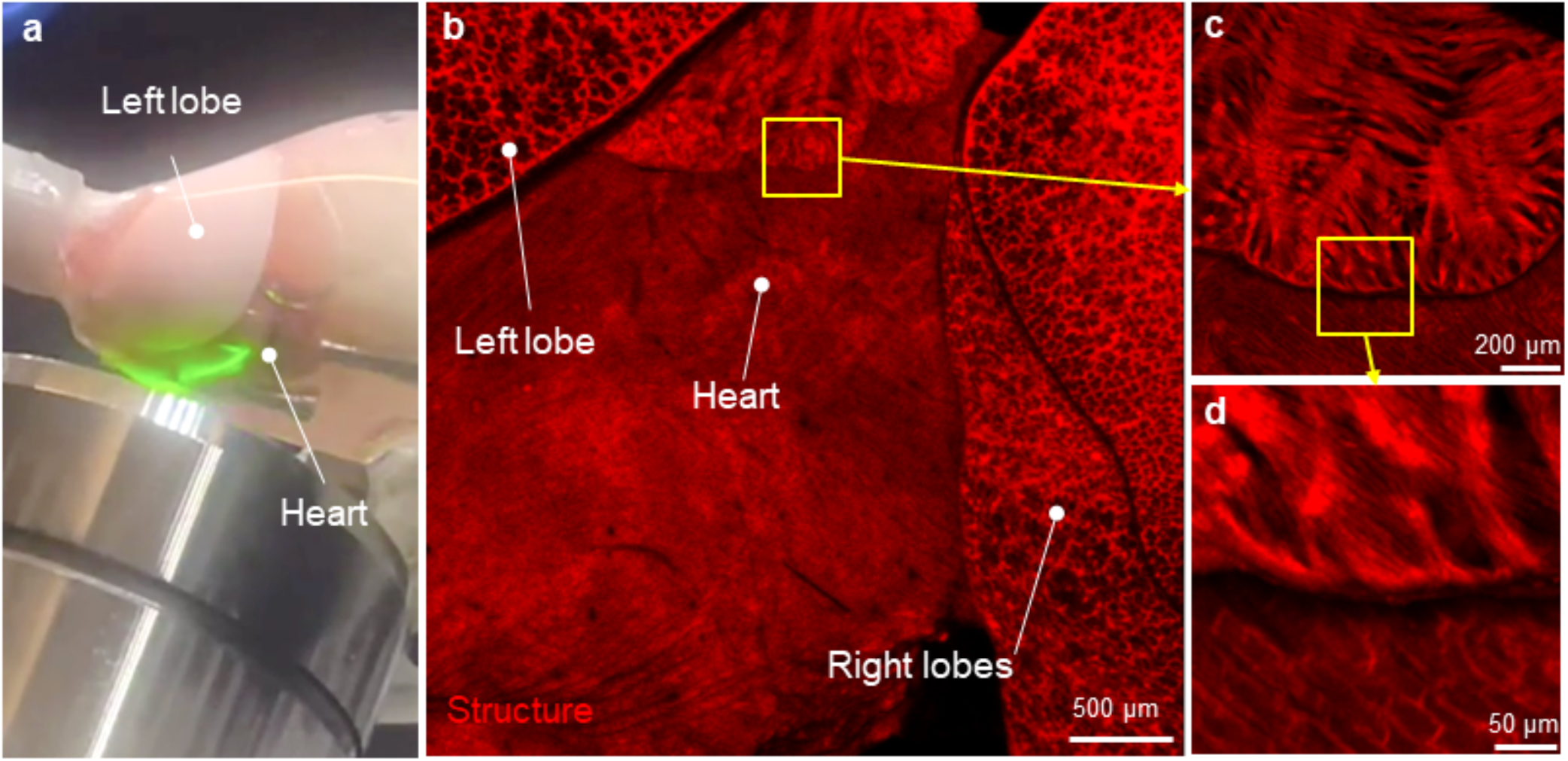
| Lung adjacent organs such as the heart can also be imaged with the crystal ribcage. Here using a laser scanning confocal (**a**) the heart is imaged with a 1.25x objective to show (**b**) the overlapping lobes of the lung followed by (**c**) higher resolution images of the atrial surface and (**d**) muscle structures.

