## Supplemental Information and Figures for "Crystal ribcage: a platform for probing real-time lung function at cellular resolution in health and disease"

Table of contents


| | | |
|---|---|---|
| **Supplemental Notes** | | |
| 1 | Crystal ribcage fabrication and extended capabilities | 47 |
| 2 | Relating mechanical alveolar strain with biological function | 47 |
| 3 | Imaging large tumor nodules on the lung | 47 |
| 4 | Alternative disruptions to respiration-circulation coupling | 47 |
| **Supplemental Methods** | | |
| 1 | Mouse ribcage 3-D object creation | 49 |
| 2 | Semi-flexible ribcage fabrication | 49 |
| 3 | Rigid ribcage fabrication | 49 |
| 4 | Crystal ribcage surface treatment | 49 |
| 5 | Versatile microscopy arm | 50 |
| 6 | Ventilation pressure and flow sensor calibration | 50 |
| 7 | Perfusion pressure and flow sensor calibration | 50 |
| 8 | Perfusion and vascular transport imaging | 50 |
| 9 | Mathematical model of diffusion | 51 |
| **Supplemental Videos list** | | |
| 1-14 | Titles of all supplementary videos | 53 |
| **Supplemental Figures** | | |
| 1 | The crystal ribcage surface is engineered to be hydrophilic, has a uniform thickness, and does not cause optical aberrations. | 55 |
| 2 | Endogenously labeled mesenchymal and immune cells imaged by confocal microscopy at whole-lobe and single cell resolution in the crystal ribcage. | 56 |
| 3 | Fibrotic regions in mice lung co-localized with increase in collagen deposition imaged using the crystal ribcage. | 57 |
| 4 | Large nodular tumors on a lung imaged through the crystal ribcage from the apex to base at alveolar resolution. | 58 |
| 5 | Tracking alveoli during changes in alveolar pressure in the crystal ribcage. | 59 |
| 6 | Image registration and strain computation workflow and validation. | 60 |
| 7 | There is no significant effect of perfusate viscosity differences between RPMI media and whole mouse blood on vascular distribution of a fluorescent tracer. | 61 |
| 8 | Nearly the entire lung surface can be imaged with the crystal ribcage. | 62 |
| 9 | Multiscale strain analysis can be performed on the alveoli and whole lung using the crystal ribcage, and capillary transport is slowest in nodular tumor and fastest in pneumonia conditions. | 63 |
| 10 | Heterogenous distribution of neutrophils (green) in an alveolar scale confocal microscopy image of LPS injured lungs. | 64 |
| 11 | Respiration-circulation coupling. | 65 |
| 12 | Local agarose blockage disrupts alveolar and vascular dynamics in the lung. | 66 |
| 13 | Air and vascular tracers leak out of lobe at site of local excision. | 67 |
| 14 | Local modulation of vascular dynamics by cauterizing the pleural lung surface. | 68 |

### Supplemental Notes

**Crystal ribcage fabrication and extended capabilities**
The crystal ribcage fabrication protocol[74] has been successfully reproduced by individuals with different training backgrounds, making it deployable to different institutions. The unique capabilities of the crystal ribcage in imaging and controllability of a functioning lung are scalable to human whole-lung, and adaptable to other organs such as heart, brain, and liver. Given the recent advances in long-term maintenance of large animal and human lung[63,83], brain[84] and liver[85], our transformative approach opens new avenues in transplantation and organ regeneration research in addition to understanding the real-time cellular pathogenesis in functional organs.

**Relating mechanical alveolar strain with biological alveolar function**
Alveolar strain is related to its true function of gas exchange in two way – (i) increasing alveolar volume during inspiration reduces gas pressure in comparison to the opening pressure of airways, driving convective gas transport in the upstream airways, and (ii) increasing alveolar size results in an increased surface area available for gas exchange across the alveolocapillary barrier[58].

**Imaging large tumor nodules in the lung**
While we are capable of studying very large nodules that replace the lung surface (**Supplemental Video 6, Supplemental Fig. 4**), we focused on smaller nodular tumors to study the local remodeling dynamics when tumors are initially established in the lung.

**Alternative disruptions to respiration-circulation coupling**
To better assess the impact of inflammation on blood flow in the capillaries, the crystal ribcage can be used with faster imaging capabilities and higher magnification power to visualize single capillary lumens (**Extended Data Fig. 8**) and future studies will utilize this capability to analyze how respiration-circulation coupling modulates red blood cell trafficking at the capillary level, and whether capillaries become blocked by the presence of immune cells during inflammation, which could cause redistribute flow to other areas. To further extend our analysis of the effect of acute injury on local respiration and circulation function, we demonstrated the capability of the crystal ribcage to probe lung functionality after surgery. We resected a small portion of tissue from the lung surface and found that the whole lobe was unable to maintain an applied alveolar pressure, as air bubbles could be seen leaking from the incision cite inside the crystal ribcage (**Supplemental Fig. 13, Supplemental Video 13).** Similarly, we labeled vascular flow through the lung with a bolus of fluorescent dye and observed that the dye began to leak out of the lung at the incision site. In the clinic, cauterization of the margins of resected lung tissue is often used to prevent air leaks and hemorrhage. We next cauterized a small portion of the lung and observed that vascular flow is faster in the cauterized region, by imaging the transport of a fluorescent dye. Additionally, the cauterized region had a shorter duration of fluorescent intensity than the surrounding tissue (**Supplemental Fig. 14, Supplemental Video 14**). This implies that cauterization does not fully eliminate vascular function but may compromise local vascular integrity such that diffusion of small molecules into the interstitium is elevated in the injured tissue. Lastly, we explored how introducing a solid phase blockage into the lung would acutely disrupt respiratory and circulatory function. We injected a small volume of fluorescently labelled agarose gel into the sub-pleura of the lung and observed that alveoli in the vicinity of the gel did not change size in response to quasi-static inflation, implying that local respiration was arrested (**Supplemental Fig. 12** and

**Supplemental Video 11**). Additionally, the distribution of a fluorescent bolus of vascular dye across the tissue surface revealed that blood flow was excluded from the gel region, indicating that, local and acute disruptions of the lung structure result in substantial loss of functionality.

#### Supplemental methods

**Mouse ribcage 3-D object creation**

Previously recorded pressure controlled μCT scans of C57B/6 and AJ mice chest cavities were obtained from the Hoffman group at the University of Iowa[75-77]. FVB mouse chest scans were performed at the Boston University Micro-CT imaging facility. These scans were segmented in MATLAB (v. R2019b Mathworks, Natick MA) using a custom algorithm and user defined regions of interest to obtain a coarse 3D object that represented the mouse chest cavity including a small section of the trachea. This coarse 3D binary object was refined and re-meshed in Meshmixer (Autodesk) to remove any sharp edges and make a stereolithography (STL) object. The Meshmixer files were imported into Solidworks 2019 (Dassault systems, France) to add registration features prior to 3D printing. The model was made age-specific by scaling the binary object by mouse lung volume reported by age[78] (**Extended Data Fig. 1**). The final STL file was printed using FormLabs (Somerville, MA) clear resin on a Form3 printer and additionally post-cured for 30 minutes. The final 3D printed insert was washed and polished to remove any layer lines and obtain a high gloss finish on the surface.

**Semi-flexible ribcage fabrication**

The 3D printed lung insert was used to make a negative mold by embedding it in a soft silicone-based elastomer called Ecoflex 00-30 (Smooth-On Inc, Macungie PA) under vacuum for 5 hours. Post curing the 3D insert was removed to obtain a smooth exact negative mold. The internal surface of the mold was coated with a thin layer of Polydimethylsiloxane (PDMS) to completely seal it from air and make the internal geometry rigid. The negative space was slowly filled with a casting sugar solution that was kept liquid at ~140˚C. The sugar filled mold was degassed in the oven for over an hour to remove any trapped air bubbles in the sugar, cured by cooling over 2 hours at room temperature, and then gently removed by deforming the soft silicone mold. PDMS was mixed in a standard 10:1 monomer to crosslinker ratio and gradually poured over the sugar mold, completely covering the surface, and then set to partially cure at ~50˚C for 25-30 minutes. This was repeated five times after which it was left to cure for 12-18 hours at ~50˚C in a dehydrating oven. The cast PDMS was positioned in a 3D printed crystal ribcage support and attached with dabs of glue. The assembly was then left in lukewarm water for 2-3 hours to dissolve the cast sugar to obtain the PDMS crystal ribcage (**Extended Data Fig. 1**).

**Rigid ribcage fabrication**

The 3D printed insert was positioned over a miniature dental thermoforming device. A 0.7mm clear polystyrene sheet was fed into the device and allowed to heat before it was pulled over the 3D printed insert while applying a vacuum to remove any trapped air between the mold and the polystyrene. The heat and vacuum were turned off and the formed polystyrene was removed from over the mold using compressed air. This rigid crystal ribcage was fitted into a support neck and fused by friction welding and using cyanoacrylate glue (**Extended Data Fig. 1**).

**Crystal ribcage surface treatment**

Both rigid and deformable crystal ribcage surfaces have native hydrophobic surfaces which were engineered to be hydrophilic. The polystyrene surface was treated with oxygen plasma using a

Harrick Plasma Cleaner, where the crystal ribcage was subjected to a vacuum of 800-900 mTorr and then plasma treatment at medium power for 2 minutes. The internal surface of the PDMS crystal ribcage was treated with a 1% PDMS-PEG Block Copolymer (DBE-712, Gelest Inc., Morrisville, PA) and allowed to cure to improve hydrophilicity as previously described [79].

**Versatile microscopy arm**

We developed two custom microscopy arms using commercially available photography brackets to allow both 3- and 6-degrees of freedom to orient the crystal ribcage with available upright (fluorescent stereomicroscope, multi-photon microscope, optical coherence tomography) or inverted microscopes (laser scanning confocal microscope) to place the desired region of interest (ROI) on the lung for imaging directly above or below the objective, and thus image any location on the lung from apex to base, on both the right and left lobes (**Extended Data Fig. 3**). For fine focusing, we use the microscopy arm in tandem with the microscope's XYZ stage controls to focus on the desired ROI with micron-scale precision. We successfully tested our mounting arm to image the lung surface with the above microscopes and with a wide range of objective power, ranging from 1.25x (lobe-scale), 10x (alveolar-scale), and 40x, and 60x (subcellular-scale) imaging of the same lung.

**Ventilation pressure and flow sensor calibration**

Ventilation pressure sensors SSDRRV100MDAA5 (Honeywell Inc.) were rated to measure between ±100 cm$H_2O$ with a 0-5 V linearly scaled output. The pressure sensor was calibrated prior to every experiment with a simple water column between 0-10 cm$H_2O$ and the calibration value was used to scale the raw voltage into pressure values in real-time. The flow sensor was calibrated one-time when it was assembled using a 3D printed constriction and SSDRRV010MDAA5 (Honeywell Inc.) differential ±10 cm$H_2O$ linearly scaled pressure sensor. The obtained voltage-flow curve which was fitted to a reflected power curve to obtain the flow values in ml/min for a given change in resistance read across the sensor. All sensor readings were scaled to real units using an Arduino and custom MATLAB functions to display in real-time.

**Perfusion pressure and flow sensor calibration**

Perfusion was measured with 26PCAFG6G (Honeywell Inc.) that were rated between 0 to 1 PSI unamplified gauge pressure sensors. The pressure sensor was read into an HX711 analog to digital converter before being read on an onboard Arduino Uno. The scaled digital values were calibrated against a known water column between 0-10 cm$H_2O$ prior to every experiment. The perfusion flow was measured using a small form factor Sensirion SLF-1300F liquid flow sensor. The sensor comes factory calibrated for DI water and 70% ethanol. It was calibrated for different perfusates by linearly scaling the sensor reading with a known syringe pump driven flow prior to the experiment. All sensor readings were scaled to real units using an Arduino and custom MATLAB functions to display them in real-time.

**Perfusion and vascular transport imaging**

Mouse lungs were perfused with serum-free RPMI at 37°C under a pressure-controlled flow of 15 cm$H_2O$, approximately 1 ml/min. To model small molecule drug transport, 50 µL of Cascade Blue dextran (10 kDa, Molecular Probes) or Cascade Blue hydrazide trisodium salt (500 Da, Invitrogen) were injected into the perfusion tubing upstream of the pulmonary artery cannula at a concentration of 10 mg/ml over approximately 1 second. Vascular dynamics were imaged over time using a laser-scanning confocal microscope (Olympus FV3000). Post-processing of vascular transport data was performed in MATLAB R2021b. Microscopy recording channels corresponding to impulse bolus injection were down sampled to a mesh grid with single pixel dimensions on the scale of a single alveolus (approximately 50 μm). Intratumor regions were segmented based on the presence of fluorescently labeled tumor cells. Within the extra-tumor regions, the peri-tumor region was segmented from regions far from the tumor based on a reduction of bolus intensity threshold. Sp3 infected areas were segmented based on the presence of fluorescently labeled neutrophils. For perfusion studies in lungs with surgical interventions, we used a microsurgical blade (AD Surgical, Sunnyvale CA) to pierce a hole approximately 250 μm into the pleural surface of the lung (**Supplemental Fig. 13**). For tissue cauterization to locally arrest ventilation, we used a small electrocautery pen (Lifeline Medical Inc., Brooksville FL) to generate burns on the pleural surface of the tissue approximately 1-2 mm in diameter. For both surgical interventions, we delivered a 50 μL bolus of Cascade Blue dextran as above to the lung during pulmonary perfusion and imaged the spatiotemporal distribution of the dye bolus in the cauterized versus uninjured tissue (**Supplemental Fig. 14**). Cauterized regions were segmented based on an increased density of red fluorescent signal due to the use of mTmG reporter mice. For studies where agarose gel was injected into the lung to acutely remodel respiration and circulation function, we loaded approximately 20 μL of 2% agarose gel prepared in water and labeled with 0.1 mg/ml Evans blue dye (ThermoFisher Scientific) into a 30G insulin syringe and injected it into the sub-pleural region of the lung, prior to placing the tissue inside the crystal ribcage. We imaged the spatiotemporal distribution of a bolus of Cascade Blue dextran in the lung vasculature during pulmonary perfusion near the gel using the same approach as above (**Supplemental Fig. 12**).

**Mathematical model of diffusion**

Fick's second law of diffusion governs the spatiotemporal dynamics of diffusion of a particular solute through a given solvent. The law states that the time derivative of the concentration of solute at a specific location is proportional to the Laplacian of its concentration at the same position.

$$(1)\ \frac{\partial \phi}{\partial t} = D\Delta\phi$$

For a ball of radius $a$ initially containing solute of constant concentration $C_0$ and bounded by a sphere perpetually maintained at constant concentration $C_1$, the concentration within the ball evolves, according to this law, as a function of radial distance and time:

$$(2)\ C(r,t) = C_1 + (C_1 - C_0)\frac{2a}{\pi r}\sum_{n=1}^{\infty}\frac{(-1)^n}{n} sin\,\frac{n\pi r}{a}\, exp\left(-\frac{Dn^2\pi^2 t}{a^2}\right)$$

where $D$ is the diffusion coefficient of the solute through the solvent. By integrating this concentration over an equatorial cross section of finite thickness, we can determine the total mass of solute within this cross section as a function of time. This leads to the expression:

$$(3)\ M(t) = \iiint_V C(r,t)dV$$

$$(4)\ M(t) = \iint_A C(r,t)LdA$$

$$(5)\ M(t) = \pi a^2 L[C_1 - 8(C_1 - C_0)\sum_{n\,\in\,\{2k-1\,|\,k\,\in\mathbb{N}\}}\frac{1}{(n\pi)^2}\, exp\,(-D(n\pi)^2 t/a^2)]$$

This last equation has a single unknown parameter, the diffusion coefficient.

In our experiments, if we assume that the intensity of the channel is directly proportional to the concentration, then we can take a unit change in intensity as our unit of concentration. It is then straightforward to compute this same quantity over a cross-section of the tumor from imaging data of the dye channel. This enables us to directly apply the previous model to the integral of the intensity of the dye channel inside the tumor.

By minimizing the squared error between the model and the dataset as a function of the diffusion coefficient, we finally recover the maximum likelihood estimate of the diffusion coefficient under the assumption of Gaussian noise.

#### Supplemental videos

Movies are in the following Google Drive folder and are included in the online submission: https://drive.google.com/drive/folders/1MAE_90I-rALiKSdqqmiD-Tqiqmbg4eIJ?usp=share_link

**Supplemental Video 1 | Functional *ex vivo* lung inside the crystal ribcage**

**Supplemental Video 2 | Whole lobe to single alveoli microscopy of lung under negative-pressure ventilation in the crystal ribcage**

**Supplemental Video 3 | Single alveoli microscopy of apex vs base in lung under negative-pressure ventilation in the crystal ribcage**

**Supplemental Video 4 | Crystal ribcage provides stable imaging surface for ventilated lung**

**Supplemental Video 5 | Immune cell migration in functional alveoli imaged in real-time through the crystal ribcage**

**Supplemental Video 6 | Apex to base image of lung with several metastatic tumors at alveolar resolution using OCT through the crystal ribcage**

**Supplemental Video 7 | Metastatic cancer cells remodel capillary function**

**Supplemental Video 8 | Real-time imaging of vascular transport at alveolar resolution in health and disease using crystal ribcage**

**Supplemental Video 9 | Neutrophil migration speed increases with higher vascular pressure**

**Supplemental Video 10 | Neutrophil extravasation and migration imaged through the crystal ribcage**

**Supplemental Video 11 | Local agarose blockage in the alveoli disrupts surrounding vascular dynamics**

**Supplemental Video 12 | Real-time imaging of spontaneous heart atrial pulsation in response to elevated pressure**

**Supplemental Video 13 | Air and perfusate leak from site of local excision in the lung**

**Supplemental Video 14 | Disrupted capillary flow in locally cauterized lung imaged through the crystal ribcage**

#### Supplemental Figures

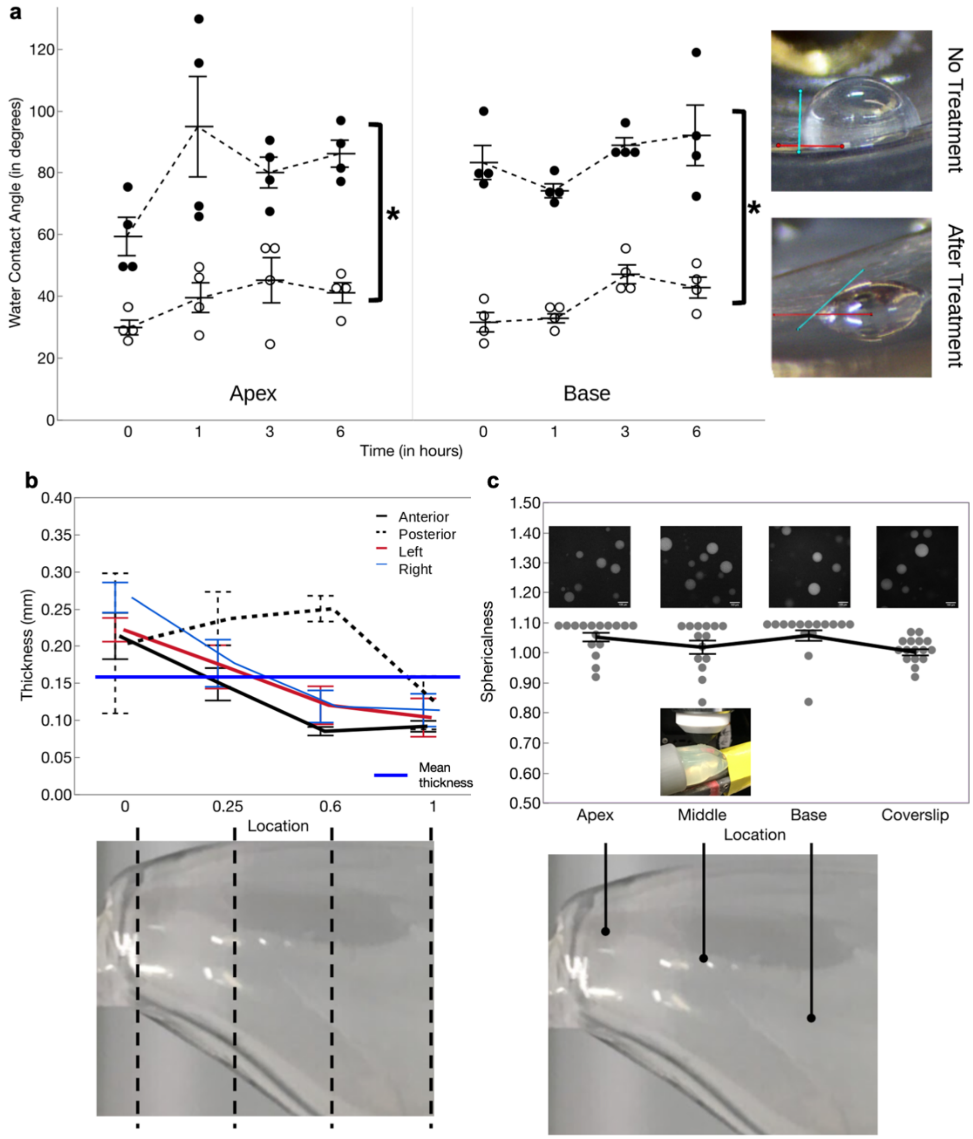


**Supplemental Figure 1 | The crystal ribcage surface is engineered to be hydrophilic, has a uniform thickness, and does not cause optical aberrations. (a)** The crystal ribcage surface retains hydrophilicity over entire surface for up to 6 hours. Hydrophilicity measured by the water contact angle of a droplet on a treated vs non-treated crystal ribcage surface. Smaller water contact angles are indicative of a hydrophilic surface. The scatter of the data from n=4 droplets at each apex and base for treated and untreated crystal ribcage is presented along with the mean ± SEM traces. **(b)** The crystal ribcage has little variation in thickness across its geometry. The crystal ribcage thickness was sampled at m=4 points at each anatomical location over n=3 shells and presented as mean ± SEM trace. **(c)** The crystal ribcage presents no significant optical aberration over the surface when compared against a coverslip. Green-fluorescent polyacrylamide beads with diameters ranging from 40-100 µm were mixed with agarose to make a slurry and uniformly spread over the surface. The beads were imaged using 2-photon microscopy. The data from m=14-16 beads at each anatomical location is presented as the mean ± SEM traces. Comparison between groups performed with two-tailed Student's t-test for significance, $p < 0.05$.

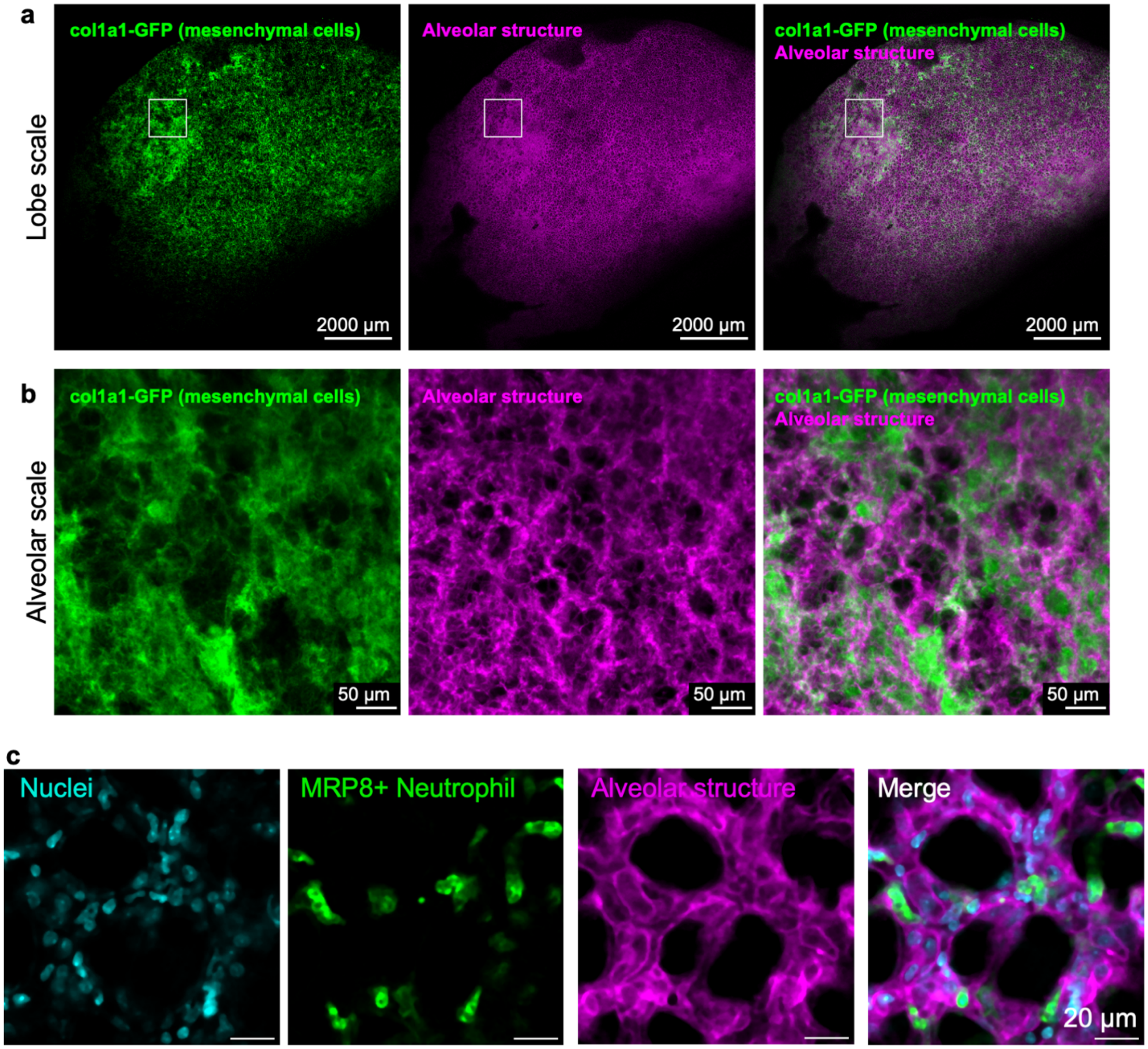


**Supplemental Figure 2 | Endogenously labeled mesenchymal and immune cells imaged by confocal microscopy at whole-lobe and single cell resolution in the crystal ribcage.** (**a**) We identified fibrotic lesions in a mouse model of pulmonary fibrosis at the lobar level through the accumulation of col1a1-GFP mesenchymal cells, achieved with col1a1 reporter mice. (**b**) col1a1-GFP$^{+}$ mesenchymal cells and lung structure remodeling can be visualized at the alveolar level in mouse models of fibrosis. (**c**) Merge of nuclei, neutrophil, and alveolar structural cells, imaged at 60x magnification with a water-immersion lens, on a laser scanning confocal microscope. Nuclei are labeled with Hoechst administered in vivo through intravascular injection. Neutrophils are labeled by membrane GFP and functional alveoli are labeled through the use of mT-MRP8-mG reporter mice.

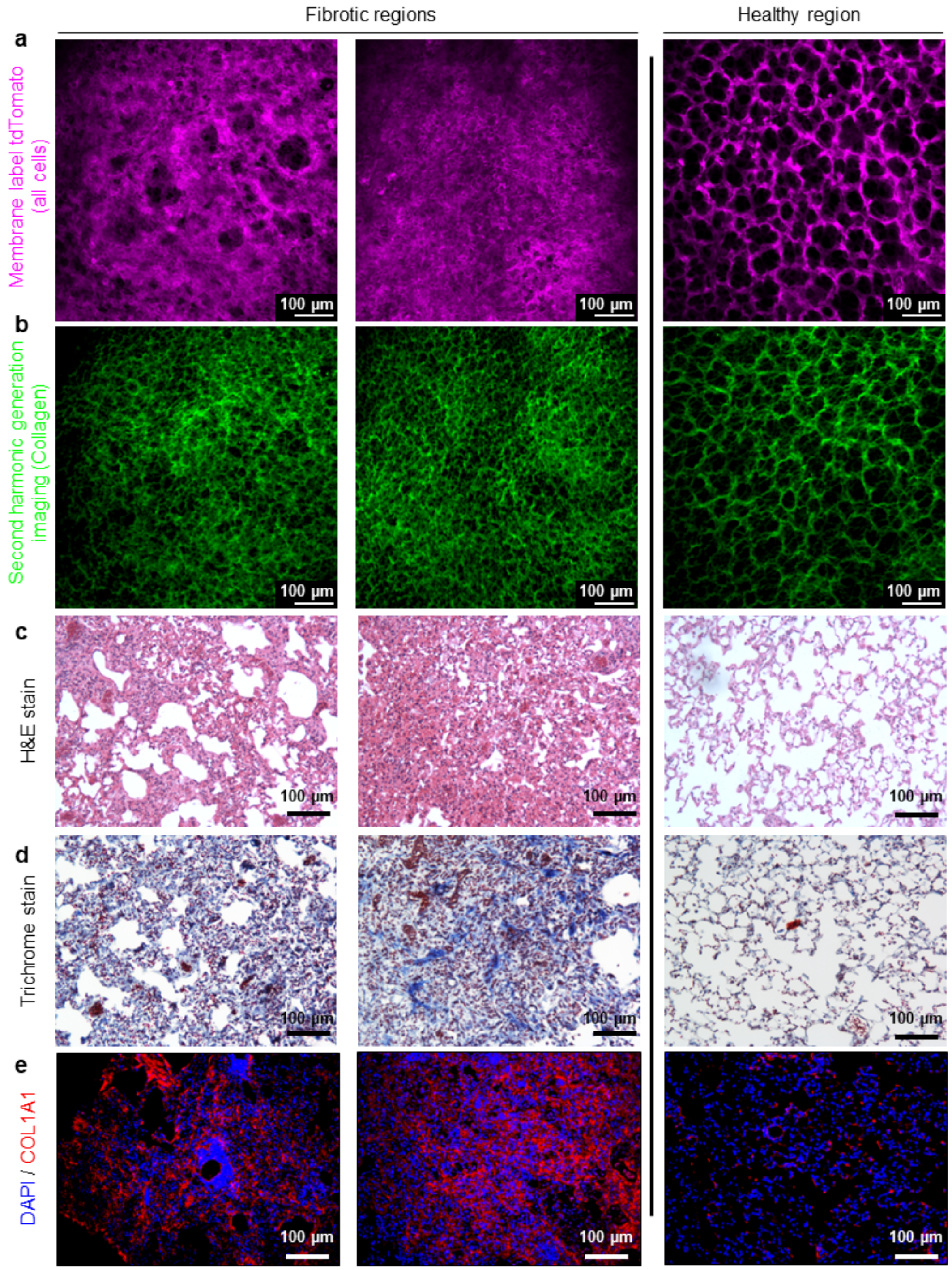


**Supplemental Figure 3 | Fibrotic regions in mice lung co-localized with increase in collagen deposition imaged using the crystal ribcage.** Both fibrotic and healthy regions imaged using two-photon microscopy to image (**a**) membrane tdTomato for all cells and (**b**) second harmonic generation (SHG) signal to see associated collagen. Healthy regions in the same lung do not have increased collagen deposition. This was confirmed against (**c**) hematoxylin and eosin (H&E), (**d**) Masson's trichrome and (**e**) Collagen-I immunofluorescence staining of the same regions that were imaged with SHG signal in the fresh lung.

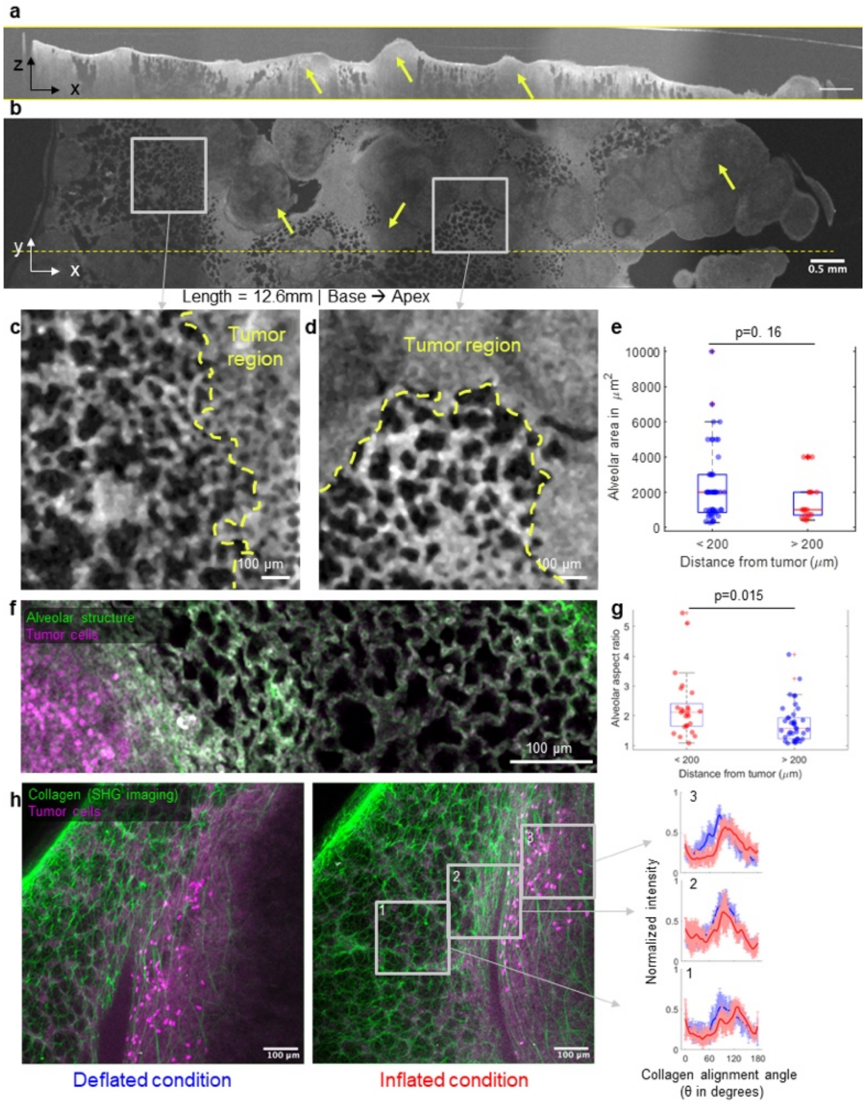


**Supplemental Figure 4 | Large nodular tumors on a lung imaged through the crystal ribcage from the apex to base at alveolar resolution.** (**a**) XZ- slice shows the bulge of nodular tumors on the lung surface (**b**) XY- slice where the total volume was stitched together from 5 sections imaged with an OCT probe. (**c, d**) Alveoli have heterogenous deformation < 200 μm from the nodular tumors and more uniform deformation >200 μm away. (**e**) n=50 alveoli show more heterogeneity in area <200 μm from nodular tumor compared to n=50 alveoli >200 μm. (**f**) Another nodule imaged in the same lung with multi-photon microscopy shows (**g**) alveoli are more elliptical with greater aspect ratio closer (< 200 μm) to the nodule for n=25 alveoli <200 μm in comparison to n=38 alveoli >200 μm from tumor. (**h**) The collagen fiber alignment (mean ± SEM) adjacent to large tumors > 1 mm (approximated from the arc of the tumor boundary in the field of view) show greater alignment when the lung is inflated. The largest tumor measured was 1.7 mm in diameter. Boxplots present median with $25^{th}$ and $75^{th}$ percentiles, whiskers are the maximum and minimum data points not considered outliers. Comparison between groups performed with two-tailed Student's t-test for significance, $p < 0.05$.

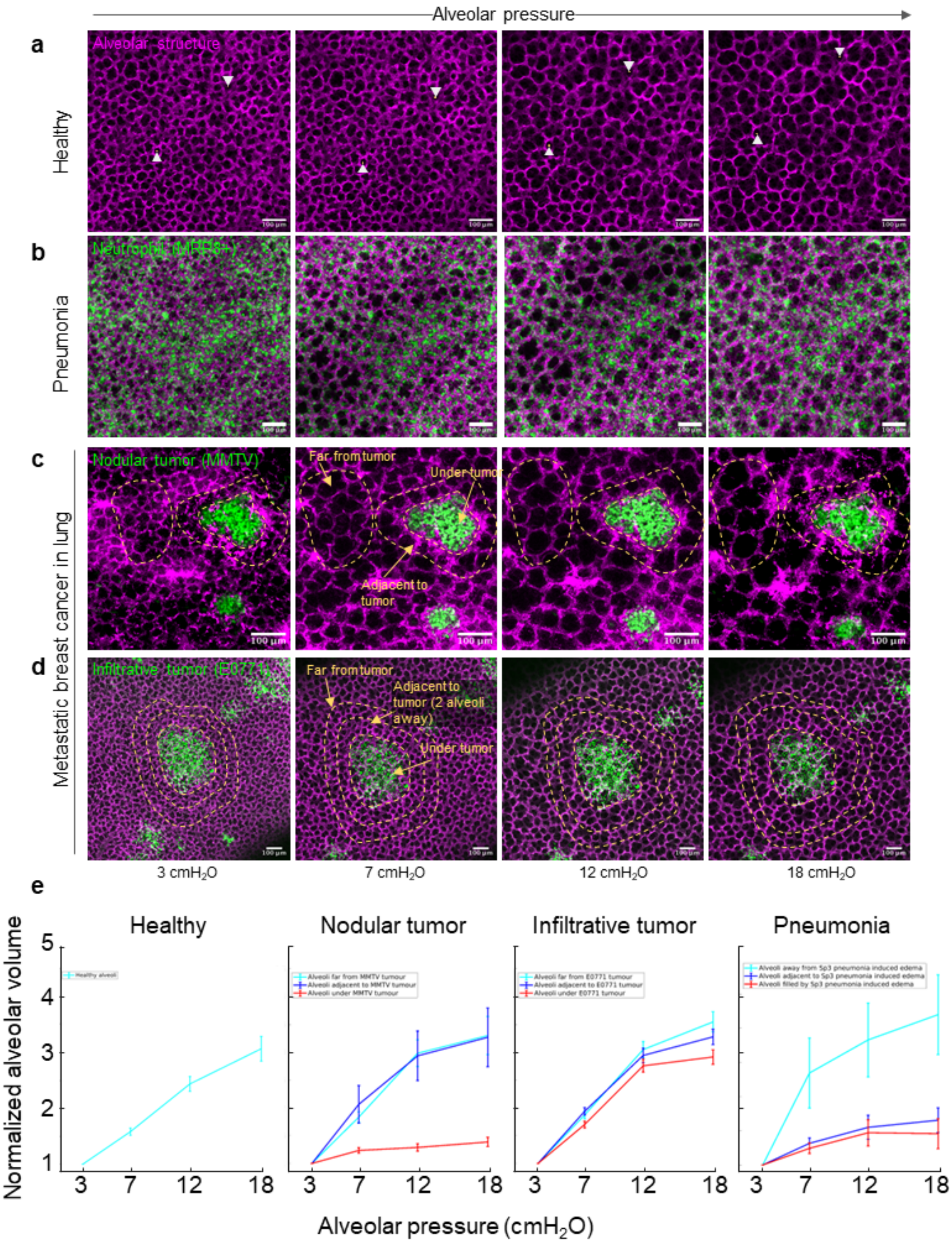


**Supplemental Figure 5 | Tracking alveoli during changes in alveolar pressure in the crystal ribcage.** (**a**) Representative confocal microscopy images for healthy, (**b**) pneumonia, and both (**c**) nodular and (**d**) infiltrative metastatic breast cancer in lung**.** (**e**) The normalized alveolar pressure-volume curve estimated from the alveolar diameter n=15 alveoli quantified for each disease condition and group at each alveolar pressure and data presented as mean ± SEM.

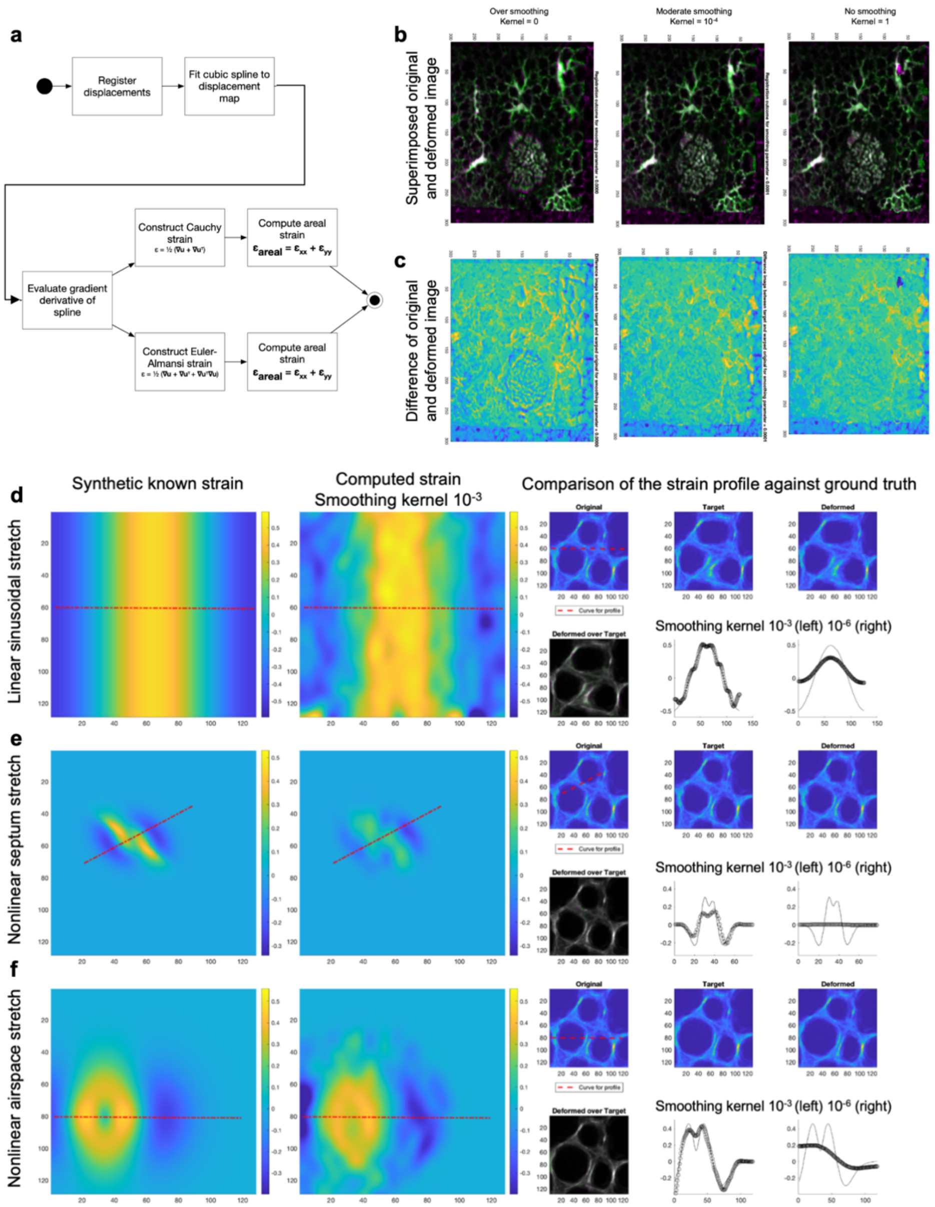


**Supplemental Figure 6 | Image registration and strain computation workflow and validation.** **(a)** Starting with image input to register the displacements, the displacement is smoothed by fitting a cubic spline which is subsequently evaluated to get a 3-D strain field from the image based 3-D displacement field. (**b**) Compared to the control / no smoothing condition with kernel = 1, a kernel = $10^{-4}$ is a moderate smoothing kernel to preserve the registration integrity while kernel = 0 over-smooths the surface (**c**) Difference image for each kernel showing variation due to smoothing. (**d**) Computed strain was validated against known linear sinusoidal stretch, (**e**) nonlinear septum stretch, (**f**) and nonlinear airspace stretch-strain fields and shows high correspondence when smoothed with a moderate kernel, here $10^{-3}$, as compared to a smaller kernel of $10^{-6}$ which does not capture the applied field.

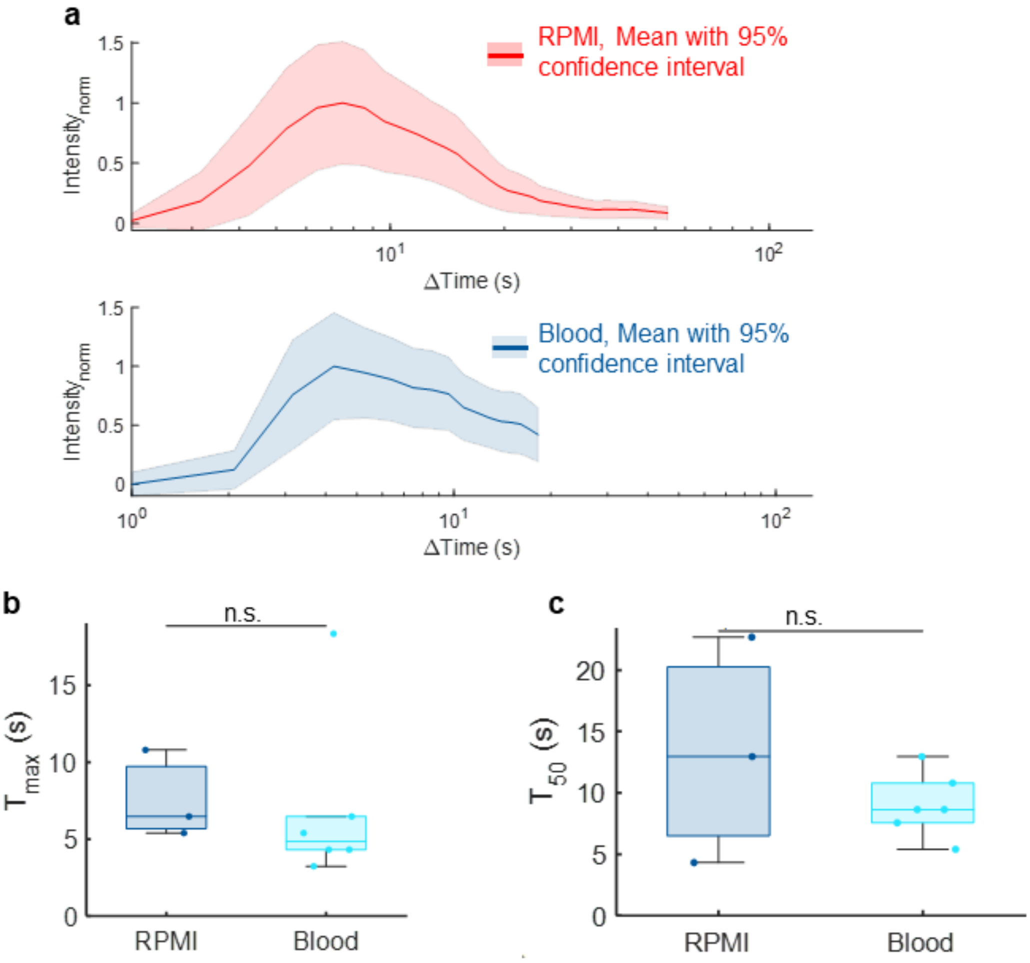


**Supplemental Figure 7 | There is no significant effect of perfusate viscosity differences between RPMI media and whole mouse blood on vascular distribution of a fluorescent tracer.** (**a**) The time-intensity distribution of the vascular dye in each RPMI and blood showed as the mean with the 95% confidence interval, (**b**) Comparison of the time at peak intensity. (**c**) The duration of the dye in the capillaries before clearing. Measurements made from m=3 ROI over n=3 mice for RPMI, and m=6 ROI over n=3 mice for whole blood. Boxplots present median with 25$^{th}$ and 75$^{th}$ percentiles, whiskers are the maximum and minimum data points not considered outliers. Comparison between groups performed with two-tailed Student's t-test for significance < 0.05.

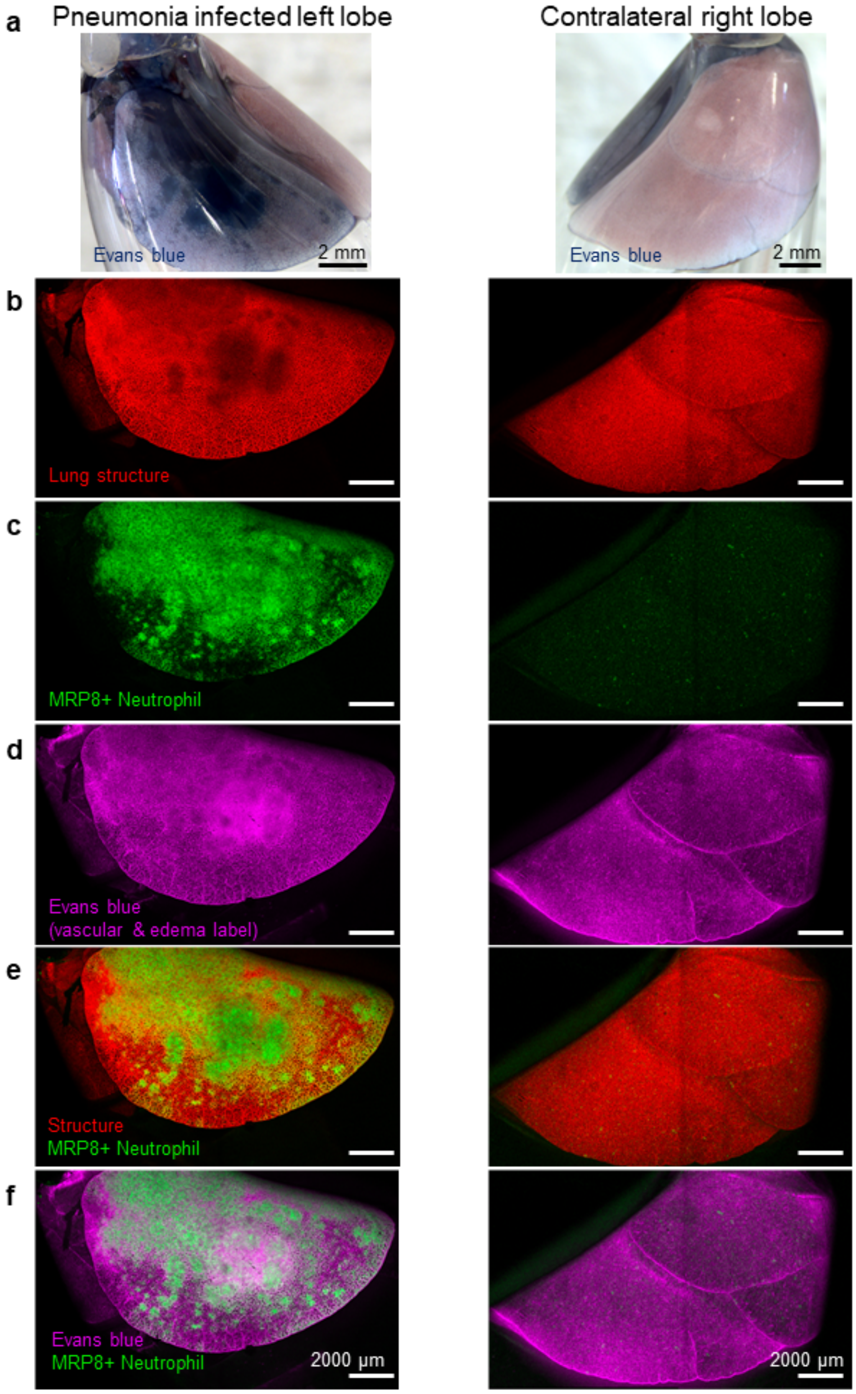


**Supplemental Figure 8 | Nearly the entire lung surface can be imaged with the crystal ribcage.** (**a**) At the whole lobe level, the pneumonia-infected left lobe and contralateral right lobe are imaged using a stereomicroscope, and each separate fluorescent marker is labeled separately for each lobe using a laser scanning confocal with 1.25x objective to get the (**b**) tdTomato labelled lung structure, (**c**) MRP8+ neutrophils, (**d**) intravascular and edema label Evans blue (**e**), combined lung structure with neutrophils (**f**), and combined edema with neutrophils. Higher resolution images of a smaller region are presented in **Extended Data Fig. 9.**

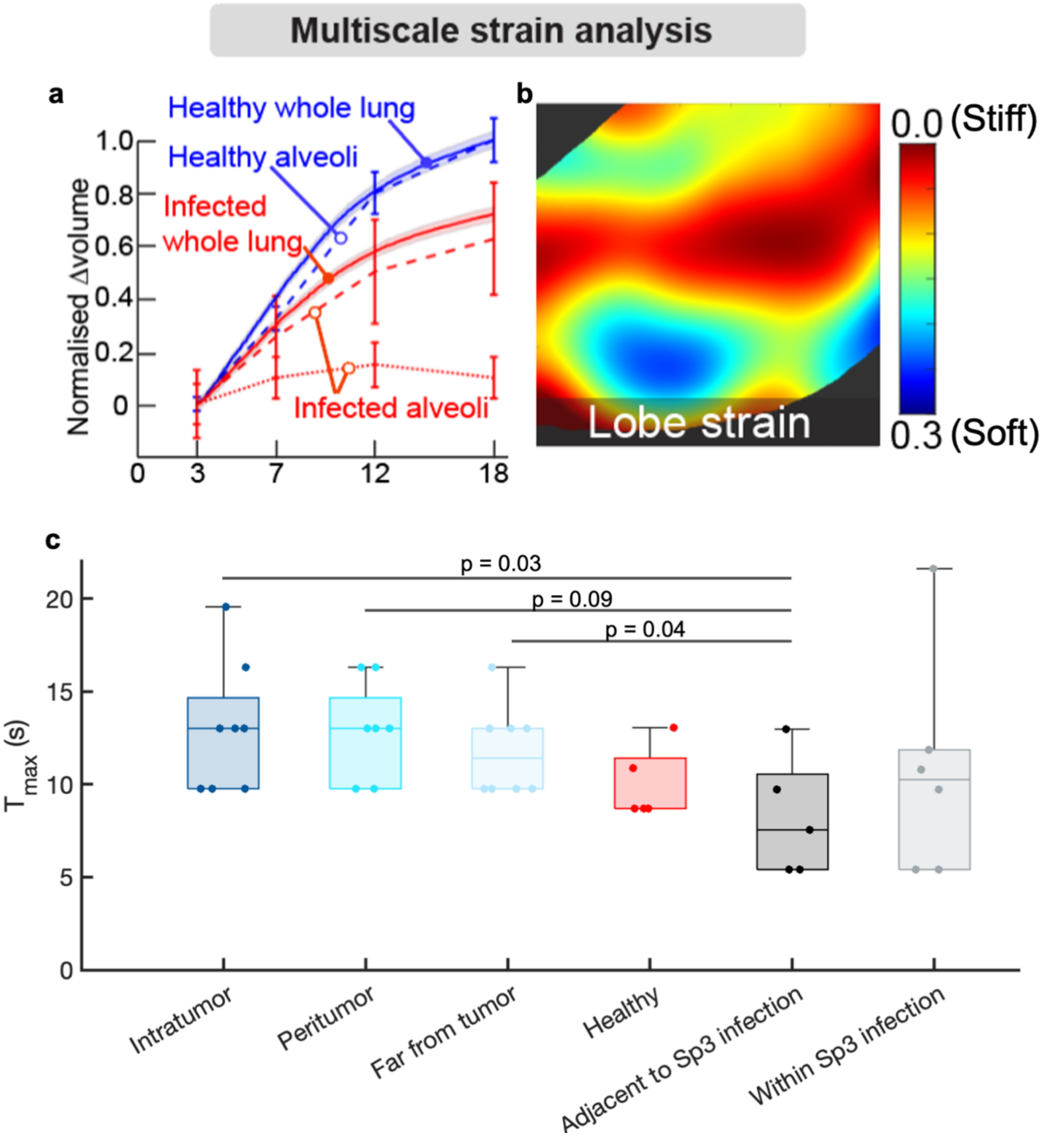


**Supplemental Figure 9 | Multiscale strain analysis can be performed on the alveoli and whole lung using the crystal ribcage, and capillary transport is slowest in nodular tumor and fastest in pneumonia conditions.** (**a**) Comparing whole lung and alveolar level pressure-volume curves (mean ± SEM) for healthy (n=3 mice) and pneumonia (n=3 mice) infected regions, whole lung pressure-volume assessment of lung function under quasi-static ventilation does not account for heterogeneity in alveolar scale function and (**b**) understanding the lobe-level strain from confocal microscopy imaging. (**c**) Time taken for dye to reach maximum intensity $T_{max}$ is least in case of pneumonia (Sp3) infection (m=9 ROI over n=4 mice) where capillaries are leaky compared to nodular tumor cases (m=8 ROI over n=4 mice) and healthy (m=4 ROI over n=5 mice), from imaging data acquired via confocal microscopy. Boxplots present median with $25^{th}$ and $75^{th}$ percentiles, whiskers are the maximum and minimum data points not considered outliers. Comparison between groups performed with two-tailed Student's t-test for significance < 0.05.

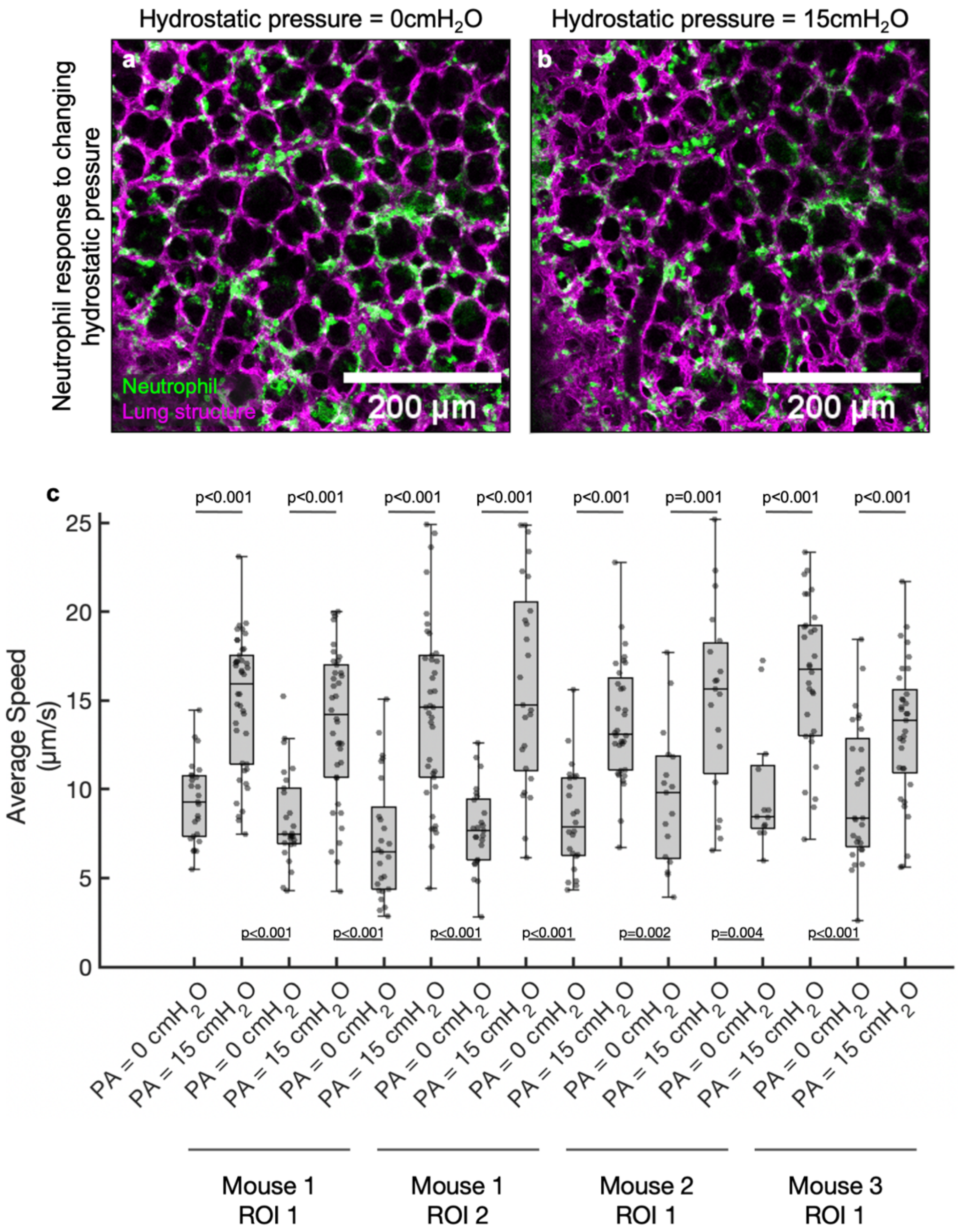


**Supplemental Figure 10 | Heterogenous distribution of neutrophils (green) in an alveolar scale confocal microscopy image of LPS injured lungs**. Alveolar pressure was held constant at 7 $cmH_2O$ while hydrostatic vascular pressure was increased from (a) 0 $cmH_2O$ to (b) 15 $cmH_2O$. (**c**) Neutrophil mechano-responsiveness demonstrated in acute lung injury (LPS). Hydrostatic pressure inside the capillaries ($P_A$) was repeated changed from 0 to 15 $cmH_2O$ in m=4 ROI from n=3 mice, from confocal microscopy image data, and neutrophil speed remained high at elevated pressure consistently. Boxplots present median with $25^{th}$ and $75^{th}$ percentiles, whiskers are the maximum and minimum data points not considered outliers. Comparison between groups performed with two-tailed Student's t-test for significance < 0.05.

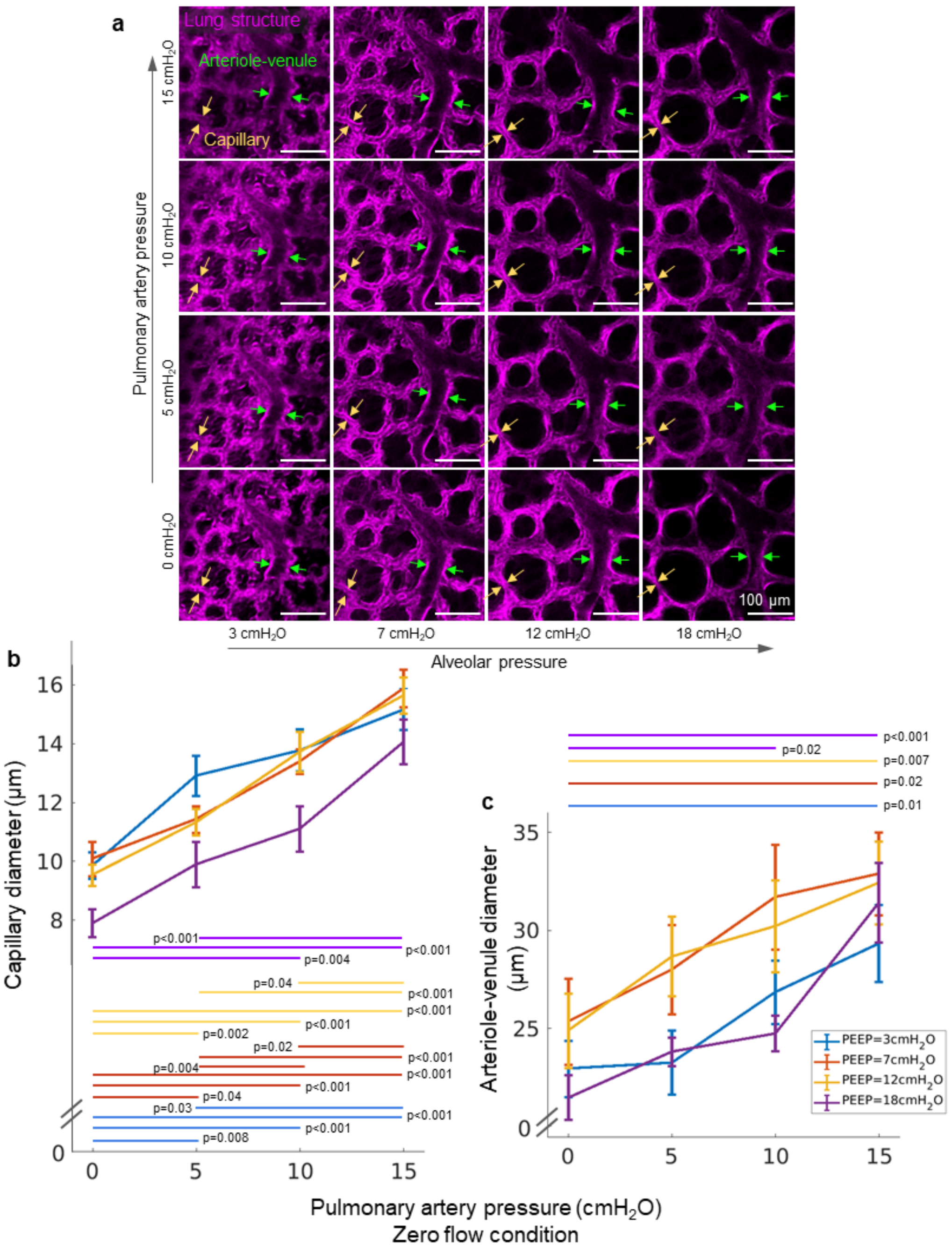


**Supplemental Figure 11 | Respiration-circulation coupling**. (**a**) Representative confocal microscopy images of arteriole-venules and capillaries imaged over the entire lung surface at different combinations of alveolar and vascular pressures, measured at the pulmonary artery, under zero-flow conditions. The changes in (**b**) capillary (m=19 ROI over n=4 mice) and (**c**) larger arteriole-venule diameter (m=15 ROI over n=3 mice). Data is reported as mean ± SEM and significance is tested with a two-tailed t-test with alpha=0.05. The vessel diameter significantly increased with increasing pressure for both micro- and medium sized vessels. However, arterioles and venules did not show a significant change in diameter when the pressure was initially increased from 0 to 5 $cmH_2O$ across all PEEP conditions.

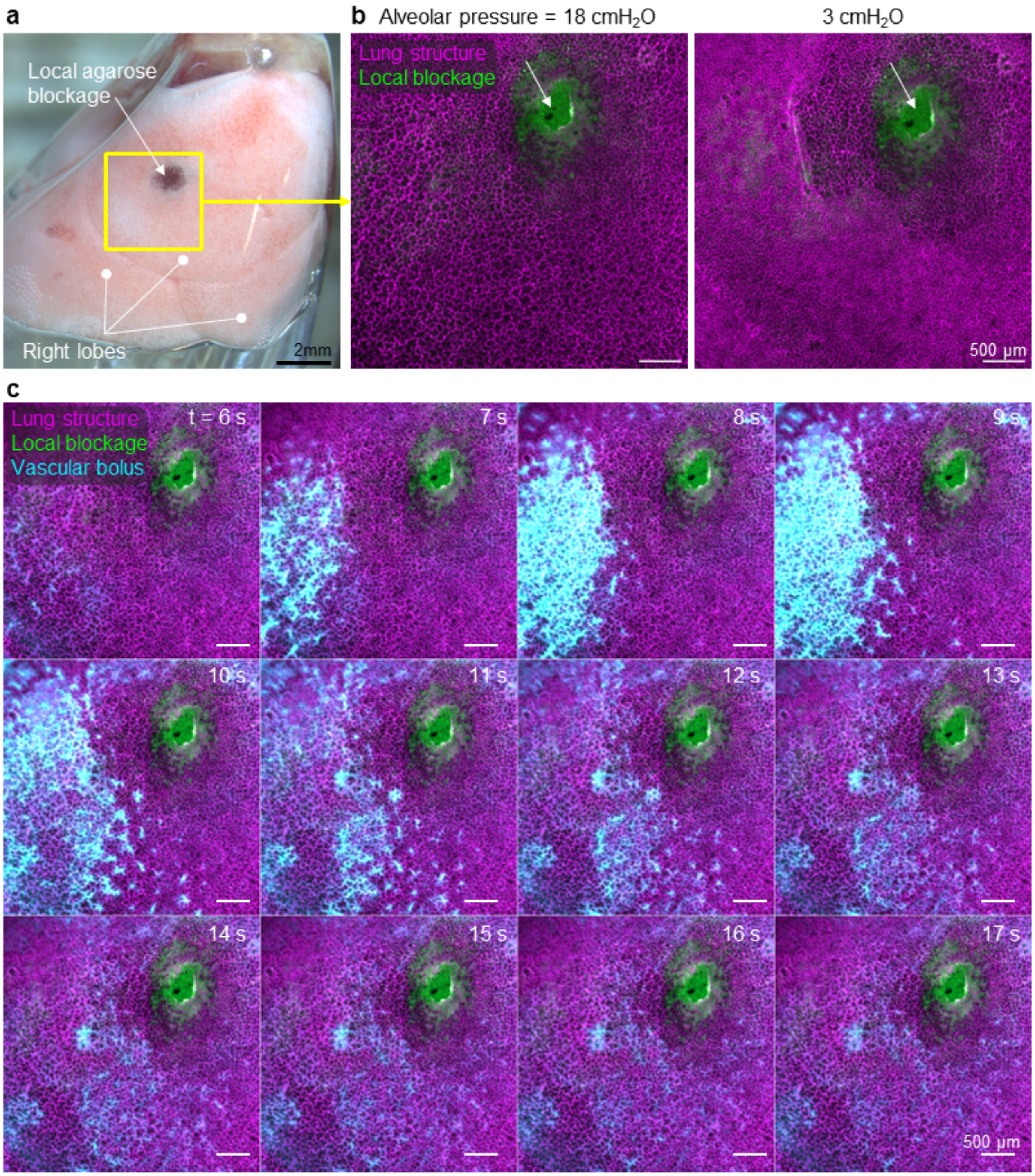


**Supplemental Figure 12 | Local agarose blockage disrupts alveolar and vascular dynamics in the lung.** (**a**) Whole-lobe stereomicroscope view showing agarose labeled with Evans blue injected into the lung subpleural space. (**b**) Confocal microscopy of alveoli near the injected agarose do not change size in response to changing air pressure at the trachea. (**c**) Timelapse imaging of the distribution of a bolus of dye perfused into the lung shows that blood flow is shunted around the injected agarose, so that the vasculature near the agarose is not perfused. Representative data shown for m=2 ROI, for n=1 mouse.

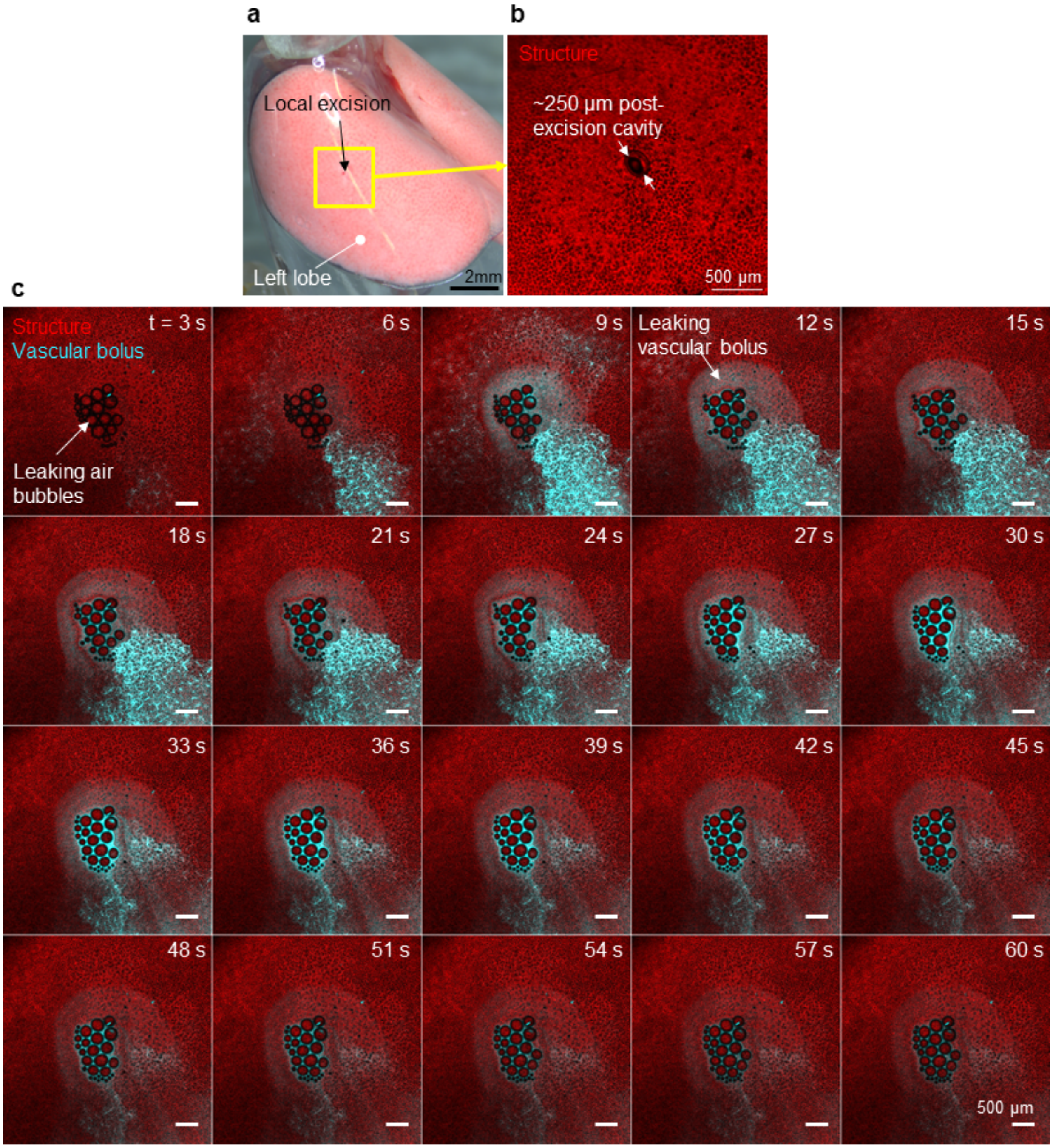


**Supplemental Figure 13 | Air and vascular tracers leak out of lobe at site of local excision.** (**a**) Stereomicroscopy of the whole-lobe view of the lung, showing a local excision of tissue on the left lobe of a healthy mouse. (**b**) Confocal microscopy view of the excision site, where lung structure is fluorescently labeled with tdTomato through using mTmG reporter mice. (**c**) Timelapse imaging of vascular transport of a bolus of dye shows that air bubbles and dye leak out of the lung at the excision site due to the compromise of alveolar and vascular integrity.

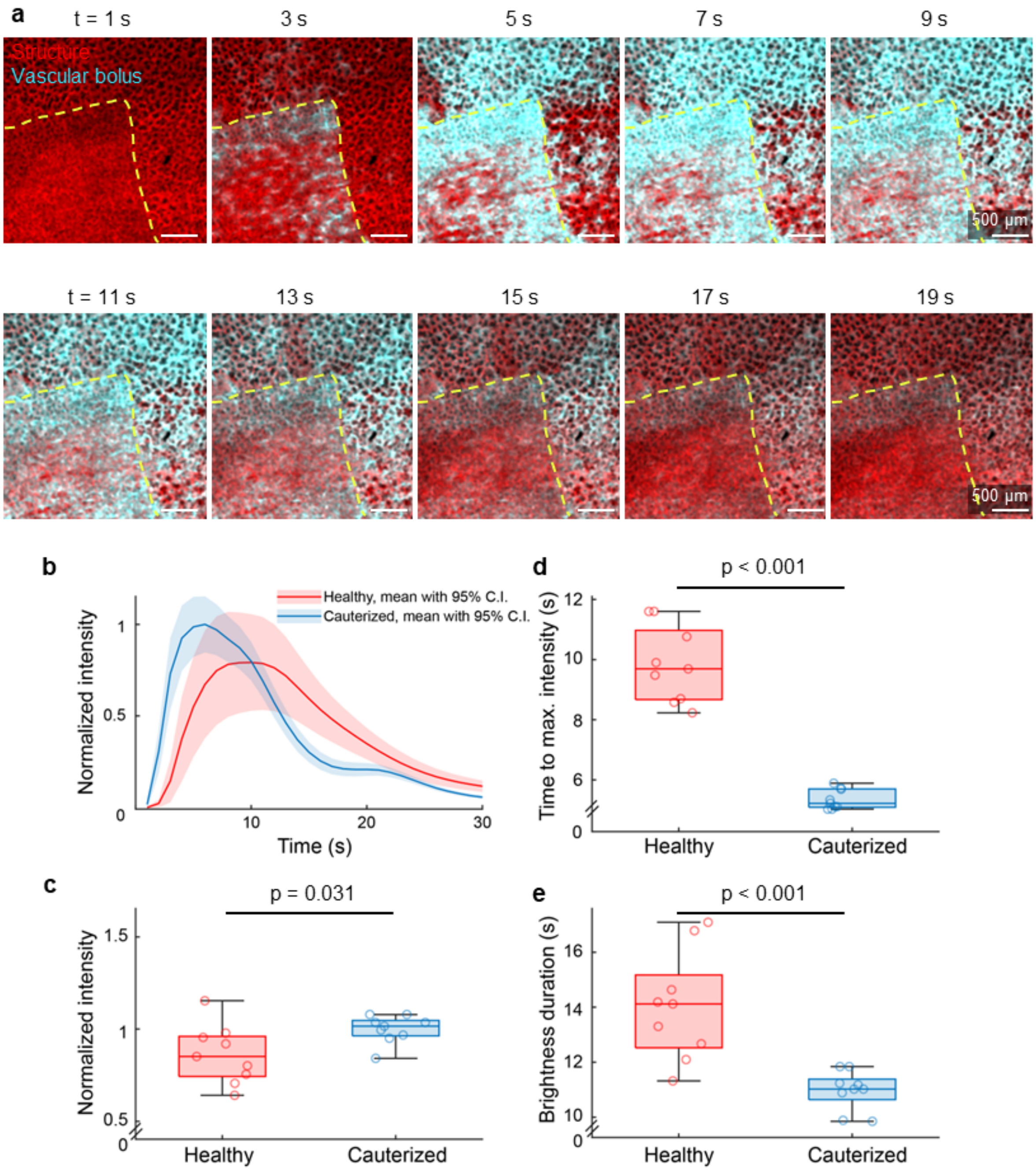


**Supplemental Figure 14 | Local modulation of vascular dynamics by cauterizing the pleural lung surface.** We used a laser scanning confocal (**a**) to image the arrival and clearance of vascular dye (10 mg/ml Cascade blue dextran) dissolved in RPMI delivered to the lung with a 1.25x objective in healthy and cauterized region. (**b**) The fluorescence intensity with respect to time for each healthy and cauterized area. (**c**) Comparison of the maximum fluorescence intensity within each region after delivery of the bolus. (**d**) Time to maximum intensity for each region after dye delivery. (**e**) Duration of brightness in each region. Boxplots show median of data and 25$^{th}$ and 75$^{th}$ quartiles as edges. Data outside these quartile ranges are indicated as hollow circles, quantified for m=9 ROI per condition from n=1 mouse.
